# Antiviral adaptive immunity and tolerance in the mosquito *Aedes aegyti*

**DOI:** 10.1101/438911

**Authors:** Michel Tassetto, Mark Kunitomi, Zachary J. Whitfield, Patrick Dolan, Irma Sánchez-Vargas, Isabel Ribiero, Taotao Chen, Ken E. Olson, Raul Andino

**Affiliations:** Department of Microbiology and Immunology, University of California, 600 16th Street, GH-S572, UCSF Box 2280, San Francisco, California 94143-2280, USA; Arthropod-borne and Infectious Diseases Laboratory, Department of Microbiology, Immunology & Pathology, Colorado State University, Fort Collins, CO 80523, USA

**Author notes:** Current address: IBM Almaden Research Center, 650 Harry Road, San Jose, California, 95120-6099, USA. These authors contributed equally to this work (Michel Tassetto, Mark Kunitomi).

## Abstract

Mosquitoes spread pathogenic arboviruses while themselves tolerate infection. We here characterize an immunity pathway providing long-term antiviral protection and define how this pathway discriminates between self and non-self. Mosquitoes use viral RNAs to create viral derived cDNAs (vDNAs) central to the antiviral response. vDNA molecules are acquired through a process of reverse-transcription and recombination directed by endogenous retrotransposons. These vDNAs are thought to integrate in the host genome as endogenous viral elements (EVEs). Sequencing of pre-integrated vDNA revealed that the acquisition process exquisitely distinguishes viral from host RNA, providing one layer of self-nonself discrimination. Importantly, we show EVE-derived piRNAs have antiviral activity and are loaded onto Piwi4 to inhibit virus replication. In a second layer of self-non-self discrimination, Piwi4 preferentially loads EVE-derived piRNAs, discriminating against transposon-targeting piRNAs. Our findings define a fundamental virus-specific immunity pathway in mosquitoes that uses EVEs as a potent and specific antiviral transgenerational mechanism.

## INTRODUCTION

*Aedes aegypti* are vectors of some of the world’s most widespread and medically concerning arboviruses such as yellow fever, dengue, Zika and chikungunya viruses. Although arboviruses can cause severe disease in humans, they generally cause non-cytopathic, persistent, lifelong infections of competent *Aedes* spp. vectors. Antiviral immunity is thought to be central to these divergent outcomes. As in other arthropods, RNA interference (RNAi) is a major component of the mosquito antiviral defense [1]. Intracellular long dsRNA, such as viral replicative intermediates generated during positive-sense viral RNA replication, are processed by the processive endoribonuclease Dicer2 (Dcr2) into 21 nucleotide (nt) small interfering RNA (siRNAs) duplexes, which are loaded onto the endoribonuclease Argonaute-2 (Ago2) at the core of the RNA-induced silencing complex (RISC). Ago2 then cleaves one strand of the virus-derived siRNA duplex and utilizes the remaining strand as a guide to target and cleave complementary viral RNAs, thereby restricting viral replication [2].

In addition to 21 nt viral siRNAs (v-siRNAs), arboviral infections of *Aedes* spp. mosquitoes and cultured cells also lead to the accumulation of 24-30 nt long virus-derived small RNAs [3][4][5][6][7][8]. These small RNAs are similar in size to PIWI-interacting small RNAs (piRNAs), which are generally associated with germline defense against mobile genetic elements. Like germline piRNAs, virus-derived piRNA (v-piRNA) production involves PIWI proteins [8][9]. In the germline, long antisense piRNA transcripts from genomic piRNA clusters are cleaved to produce primary piRNAs with a uridine at their 5’ end (U1), which are loaded onto a first Piwi protein (Piwi or Aubergine in *Drosophila* [10]) to target and cut cognate transposon mRNAs. The 3’ products from cleaved transposon mRNAs thus bear an adenine at their new position 10 (A10), complementary to the U1 of antisense piRNAs and are bound by Ago3. The new piRNA complex can then target its corresponding antisense transcript to generate a new antisense U1 piRNA. This ping-pong model thereby provides the germ line with a secondary biogenesis pathway of anti-transposon piRNAs. Although, v-piRNAs are predominantly of the same orientation (sense) than the viral genome/mRNA, antisense v-piRNAs are also produced and provide evidence of ping-pong amplification of v-piRNAs in alphavirus-infected mosquitoes [6][10]. However, v-piRNAs are not observed in virus-infected Drosophila, nor are PIWI proteins involved in the fly antiviral defense[11]. Despite a much greater invasion of its genome by TEs compared to the fruitfly, the *Aedes* mosquito dedicates a smaller fraction of its piRNAs to target TEs [12] and possess an expanded Piwi gene family compared to *Drosophila* [13], suggesting a functional diversification of the Piwi family in mosquito. Conflicting reports have attributed antiviral roles to different PIWI clade proteins in mosquito. For instance, Piwi5 and Ago3 have been suggested to be specifically required for the formation of virus-derived piRNAs, though knock-down of these proteins had no effect on viral replication [8] [14]. By contrast Piwi4, has been shown to be antiviral in dsRNA knock-down screens [14] [9] but a direct association of viral piRNAs with Piwi4 has not been shown yet [8].

Upon arboviral infection in insect cells, fragments of viral RNA genomes are also converted into vDNA by the reverse-transcriptase activity of endogenous retrotransposons, leading to the accumulation of episomal vDNA molecules. These episomal vDNA molecules serve in the amplification of the antiviral RNAi response in Drosophila [15] and have been linked to vector competence in mosquito[7]. At the genomic level, integration of viral elements has been identified in all eukaryotes and for all virus families[16]. In insects, these endogenous viral elements (EVEs) derived mostly from non-retroviral RNA viruses[17][18], with *Aedes spp* genomes harboring approximately 10 times more EVEs than other mosquito species. Parallel analyses of genome and small RNA sequences from *Aedes* mosquitoes and derived cell lines reveal that EVEs organize in large loci characterized by high LTR retrotransposon density and the production of high levels of piRNAs[18]. The abundance of EVEs and EVE-derived piRNAs in mosquito cells, the accumulation v-piRNAs during infection and the contribution of PIWI proteins in antiviral defense suggest the existence of EVE-derived immunity in mosquito through RNAi mechanisms. However, from a mechanistic point of view, little is known about the role of the piRNA machinery in the RNAi-based adaptive immunity in mosquito and the potential antiviral function of EVEs and.

Here we demonstrate that Piwi4 is upregulated in somatic tissues in adult female *Ae. Aegypti* following blood meal and restricts dengue virus replication *in vivo*. In infected cells, Piwi4 is required for the accumulation of mature v-piRNAs and binds preferentially to antisense v-piRNAs (corresponding to the anti-genome viral RNA) and not to anti-transposon piRNAs. High-throughput sequencing of episomal DNA accumulating during acute arboviral infection reveals that unlike host mRNAs, viral RNAs are preferentially reverse-transcribed by retrotransposons, indicating that this antiviral system discriminates between self and non-self RNAs during the vDNA acquisition process. In the context of the persistent Cell Fusing Agent virus (CFAV) infection in Aag2 cells, we show that Piwi4 preferentially binds to v-piRNAs derived from genomic CFAV EVEs and not from the replicating CFAV RNA. Conversely, knock-down of Piwi4 results in a decrease in CFAV EVE-derived piRNAs and an increase of CFAV RNA accumulation. Furthermore, Aag2 cells infected with Sindbis virus engineered to carry EVE sequences lead to efficient inhibition of SINV replication in a sequence and strand-specific manner. Together our results indicate that genomic EVEs, EVE-derived piRNAs and Piwi4 are central to the antiviral adaptive immunity in *Ae. aegypti* mosquitoes. Given that EVEs are stably incorporated in to genome, these observation illuminate the molecular basis for the acquisition of transgenerational RNAi immunity.

## RESULTS

### Piwi4 is required to restrict virus replication in mosquitoes

In *Ae. aegypti*, four Piwi clade proteins have been associated with the host response to arboviral infection. While Piwi5 and Ago3 are involved in v-piRNA production, their knock down does not affect viral replication [19] [8]. By contrast, knock-down of Piwi4 expression leads to increased replication of Semliki Forest Virus, Bunyawera virus, Zika virus and Rift Valley fever virus but does not affect the overall amount of v-piRNAs [9] [20] [21]. Therefore, we first confirmed the role of PIWI proteins in antiviral defense using a RNAi screen against all seven Aedes Piwi transcripts, followed by Sindbis virus infection (SINV; *Alphavirus*; *Togaviridae*). We found that Piwi4 had the strongest antiviral effect in Aag2 cells (Figure 1 - supplement A). Similarly, Piwi4 knock-down in Aag2 cells led to higher viral replication of dengue virus type 2 (DENV2; *Flavivirus; Flaviviridae*) and Chikungunya virus (CHIKV; *Alphavirus*) (Figure 1A).

All previous studies of Piwi proteins in mosquito antiviral defense have been conducted in cell culture. To relate these findings to the actual insect vector, we examined Piwi4 expression in somatic tissues of adult female *Ae. aegypti* mosquitoes and observed transient increases in both midguts and carcasses, but not in the ovaries, after a blood meal (Figure 1B, *p* ≤ 0.01-0.05, Mann-Whitney U test). The antiviral role of Piwi4 in female *Ae. aegypti* was then tested by anti-Piwi4 dsRNA injection (dsPiwi4) followed by infection with DENV-2 (Jamaica 1409) by blood meal. Depletion of Piwi4 (dsPiwi4, Figure 1 - supplement B) resulted in a significant increase of DENV2 genomic RNA in both the midgut and carcass at 7 and 10 days post infection (dpi) and a significant increase of infectious virus titers in whole mosquitoes at 10 dpi (Figure 1C, *p* ≤ 0.05, Mann-Whitney U test). These results indicate that Piwi4 participates in antiviral defense not only in cultured Aag2 cells but also in *Ae. aegypti* somatic tissues.

**Figure 1.**
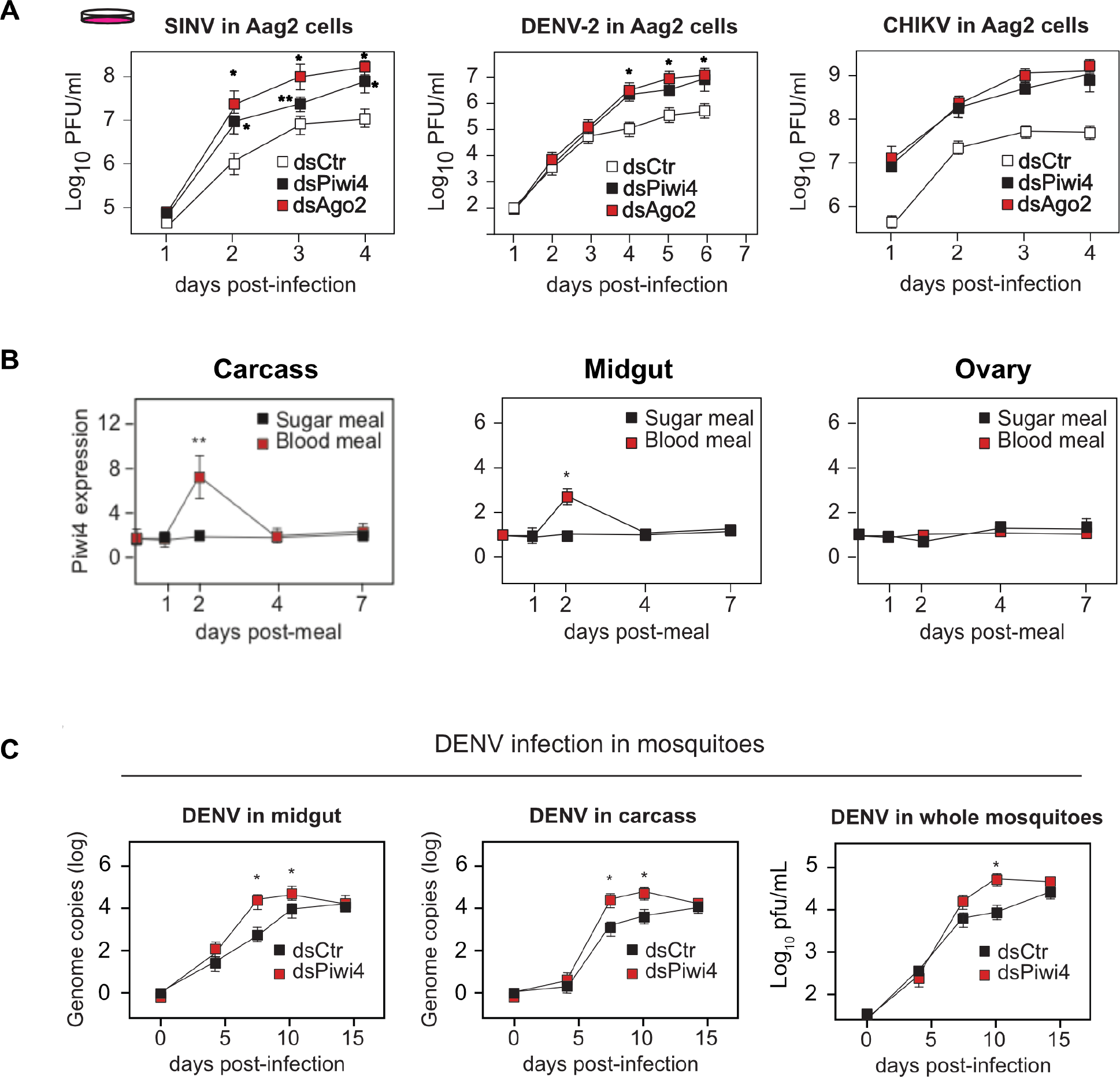

### Piwi4 is required for v-piRNA maturation

What are the mechanisms of Piwi4 antiviral activity? Although Piwi4 has not been reported to affect overall v-piRNAs accumulation, its role in v-piRNA maturation has not been thoroughly investigated. To re-assess the role of Piwi4 in the production of mature antiviral small RNAs, we analyzed virus-derived small RNAs from Ago2, Piwi4 or Ago3 depleted Aag2 cells, infected with SINV (Figure 2 - supplement A). In control dsRNA treated cells, both SINV-derived siRNAs (v-siRNAs, 21 nt long) and SINV-derived piRNAs (v-piRNAs, 24-30 nt long) accumulated during infection (Figure 2A). The 21 nt v-siRNAs were uniformly derived from the entire viral genome, representing both the sense and antisense strands of the genome (Figure 2 - supplement B). In contrast, 24 to 30 nt v-piRNAs were asymmetrically distributed with respect to strand polarity and position on the virus genome (Figure 2 - supplement C). As previously described[8][9], the overall size distribution of v-piRNAs was not affected by Piwi4 knock down. We then asked whether v-piRNA maturation was affected by Piwi4 loss-of-function. Maturation of piRNAs is mediated by their 3’-end methylation following loading onto a PIWI protein and can be readily assessed by its resistance to ß-elimination[22]. Analysis of virus-derived small RNAs after ß-elimination showed a decrease in mature v-piRNAs in Piwi4 knock-down samples (Figure 2B), suggesting a role for Piwi4 in v-piRNA maturation.

**Figure 2.**
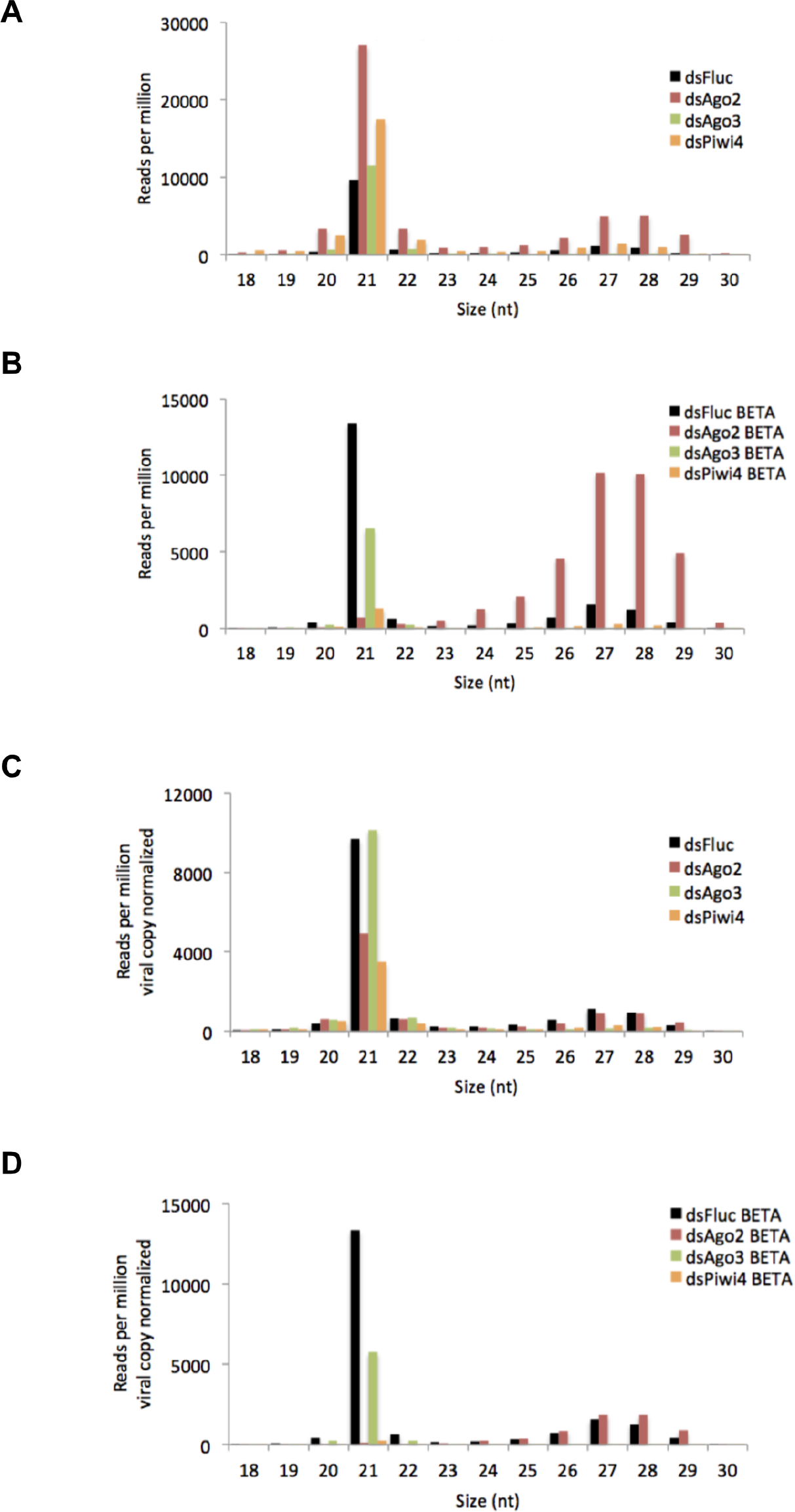

The large accumulation of sense v-piRNAs from the 5’-end of SINV subgenomic RNA (genome nt 7611-11703) is consistent with a model in which piRNA ping-pong amplification correlates with the abundance of RNA target (for instance, SINV subgenomic RNA) (Figure 2 - supplement C). To accurately assess the effect of gene-knockdown on v-siRNAs and v-piRNAs maturation and avoid biases due to differences in viral RNA abundance between conditions (Figure 2 - supplement D), read counts were normalized to SINV genome copy numbers (Figure 2C). Knockdown of either Piwi4 or Ago3 significantly reduced the abundance of mature v-piRNAs (resistant to ß-elimination, Figure 2D) and transposon-derived piRNAs (Figure 2 - supplement E and F). However, knockdown of Piwi4 or Ago2, but not Ago3, resulted in complete depletion of mature v-siRNAs (Figure 2D). Our results indicate that Piwi4 is involved in v-piRNA and v-siRNA maturation but appears to make a minor contribution to their initial production (see Discussion).

### Piwi4 binds to antisense v-piRNAs

Because maturation of piRNAs is linked to their association with PIWI proteins [22], we then assessed whether Piwi4 could bind to v-piRNAs. To address this, we performed Piwi4-immunoprecipitation (IP), using a Piwi4-FLAG construct transiently expressed in SINV infected Aag2 cells, followed by small RNA sequencing. To control for background resulting from the high abundance of small RNAs in the cells and potential confounding effects due to Piwi protein overexpression, we normalized small RNAs associated with Piwi4 to small RNAs present in the input sample. Contrary to previous reports that did not normalize to input small RNA content, we found that Piwi4 bound to v-piRNAs and that Piwi4-associated small RNAs were significantly enriched for antisense SINV-derived piRNAs (Figure 3A, antisense vs sense v-piRNAs, *X*^*2*^, *p* < 2.2e-16). In addition, Piwi4-associated v-piRNAs displayed a uracil bias at their first position (Figure 3B), a signature of Piwi-bound piRNAs. In contrast, Piwi4 did not significantly associate with v-siRNAs (Z score for 21 nt sense and antisense v-siRNAs enrichment −0.2 and −0.6 respectively) nor TE-derived piRNAs (Figure 3C).

**Figure 3.**
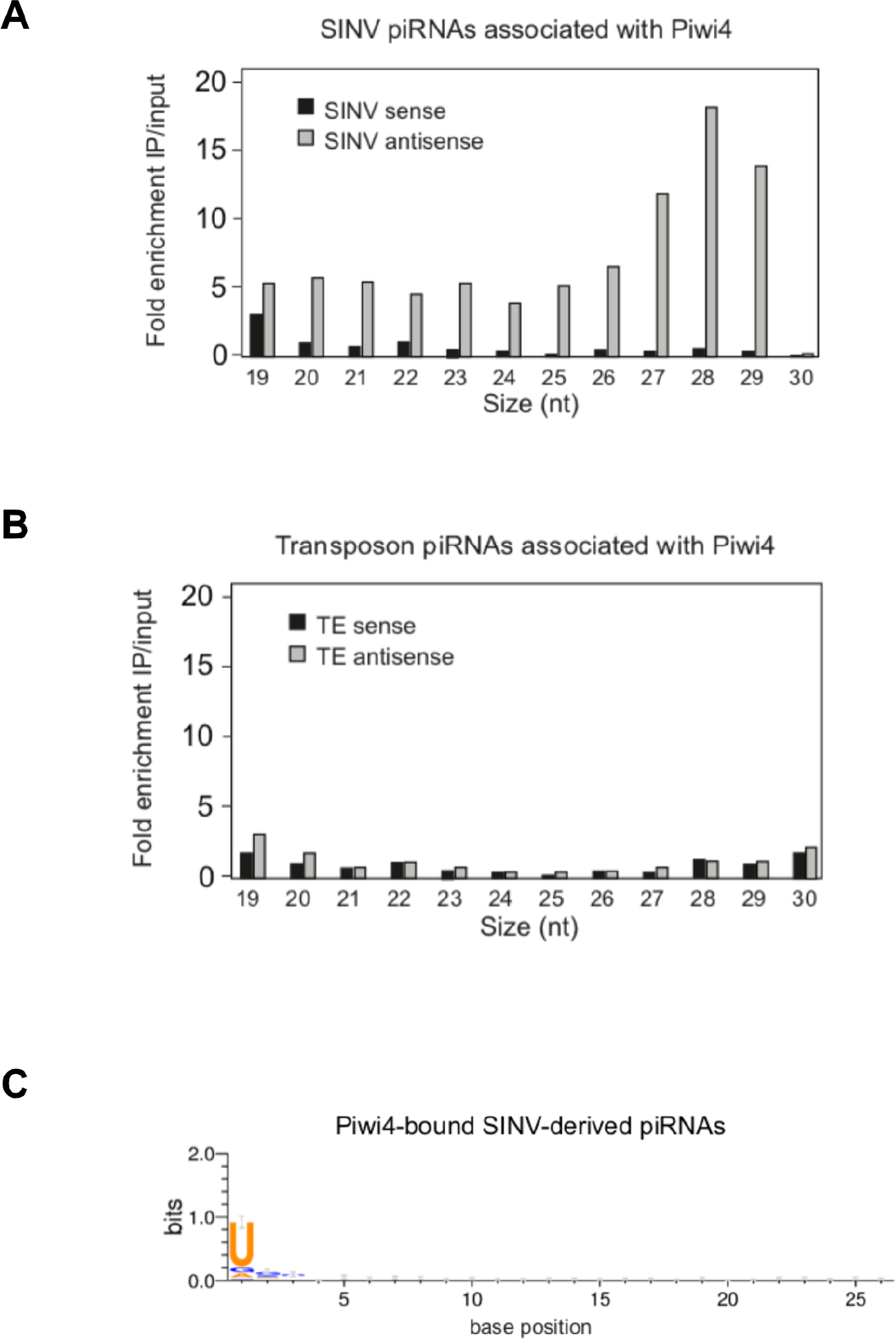

To further validate that Piwi4 preferentially bound to antisense v-piRNAs, we performed three additional IP followed by small RNA sequencing experiments in SINV infected Aag2 cells: two using the Piwi4-FLAG construct and one negative control IP experiment in Aag2 cells transiently expressing eGFP. Deep-sequencing analysis of small RNAs in Piwi4-IP and input fractions confirmed that Piwi4 preferentially binds to antisense v-piRNAs (antisense vs sense v-piRNAs, *X*^*2*^, *p* < 2.2 e-16 and *p* < 4.2e-9, for Experiment 1 and 2 respectively) with a U1 bias, during acute SINV infection in Aag2 cells (Figure 3 - supplement A). In addition, v-piRNAs enrichment in Piwi4 IP experiments were significantly greater than in the eGFP control experiment (Figure 3 - supplement B, *X*^*2*^, *p* < 1.497e-12 and *p* < 2.2e-16 for 26-29 nt-long antisense v-piRNA enrichment in eGFP vs Exp1 and Exp2 respectively). Of note, normalization to input small RNA content revealed a lack of v-siRNA enrichment in the Piwi4-bound fraction, consistent with the size preference of PIWI proteins [10]. This suggests that the large amount of v-siRNAs accumulating during SINV infection (Figure 2A and [8]) was non-specifically immunoprecipitated with Piwi4.

These results further substantiate the idea that Piwi4 is functionally distinct from other Piwi proteins involved in defense against TE and viruses. In agreement with a recent study[9], we found that Piwi4 interacts with Ago2 (Figure 3 - supplement C). Because knock-down of Piwi4 expression affects v-siRNAs production (Figure 2D) but does not specifically bind to them (Figure 3A and supplement), our results suggest that the siRNA and piRNA pathways are linked through Piwi4, which may act as a hub to coordinate production of antiviral small RNAs.

### Aedes aegypti cells conversion of viral RNA genome into vDNA efficiently discriminates between viral and host RNAs

During infection of *Drosophila melanogaster* [15] and in mosquitoes [7][23][24], fragments of RNA virus genomes are reverse-transcribed early during infection, which results in viral cDNA formation (vDNA). Expanding on these studies that used the reverse transcriptase (RT) inhibitors AZT to block vDNA formation, we found that treatment of SINV-infected Aag2 cells with another RT inhibitor, Stavudine/d4T, also led to elimination of SINV-specific vDNA (Figure 4 - supplement B) and higher SINV replication in Aag2 cells (Figure 4A). Also, AZT and d4T treatments resulted in a significant reduction of v-piRNA reads but caused increased production of v-siRNAs (Figure 4B and Figure 4 - supplement A). These results provide a direct link between reverse transcription of viral genomes, production of piRNAs and the antiviral response in mosquito cells. Our data, together with previous observations[23][7], suggest that during acute arbovirus infection of mosquito somatic tissues, primary v-piRNA precursors are transcribed from *de novo* synthesized vDNA, which are then processed into mature antiviral v-piRNA bound preferentially by Piwi4 to repress viral replication (Figure 4C). In plants and animals, retrotransposons produce circular DNA intermediates as part of their replication cycle[25][26][27]. We, and others, have hypothesized that vDNA in infected mosquito cells is produced in the context of retrotransposon replication (Figure 4C). In support of this model, we observed that PCR amplification of circular episomal DNA prepared from SINV-infected Aag2 cells with SINV genome-specific primers yielded a greater amplicon concentration than SINV-specific amplification of genomic DNA prep (Figure 5 - supplement A). To directly analyze the role of retrotransposons in vDNA formation, we infected Aag2 cells with SINV and purified cytoplasmic extra-chromosomal DNA enriched for circular DNA[28], followed by Nextera cloning and Illumina deep-sequencing (Figure 5A). In addition to sequences derived from the circular mitochondrial genome (1% of total reads), we found that our preparations contained a significant enrichment of reads mapping to transposons (10-22% of total reads). Transposon sequences came almost exclusively from retrotransposons, as expected due to a cytoplasmic step in their replication cycle. LTR-retrotransposons (Ty1, Ty3 and Pao Bel) were the most abundant sub-class observed (Figure 5B). Importantly, we also observed a significant number of reads in the circular DNA enriched fraction mapping throughout most of the SINV genome (Figure 5D, *i*). Given that Aag2 cells are persistently infected with the insect-specific flavivirus, cell fusing agent virus (CFAV)[29], we reasoned that the circular DNA preparation should also contain vDNA derived from the CFAV genome. Indeed, we observed a large number of reads mapping uniformly throughout the CFAV genome (Figure 5D, *ii*).

**Figure 4.**
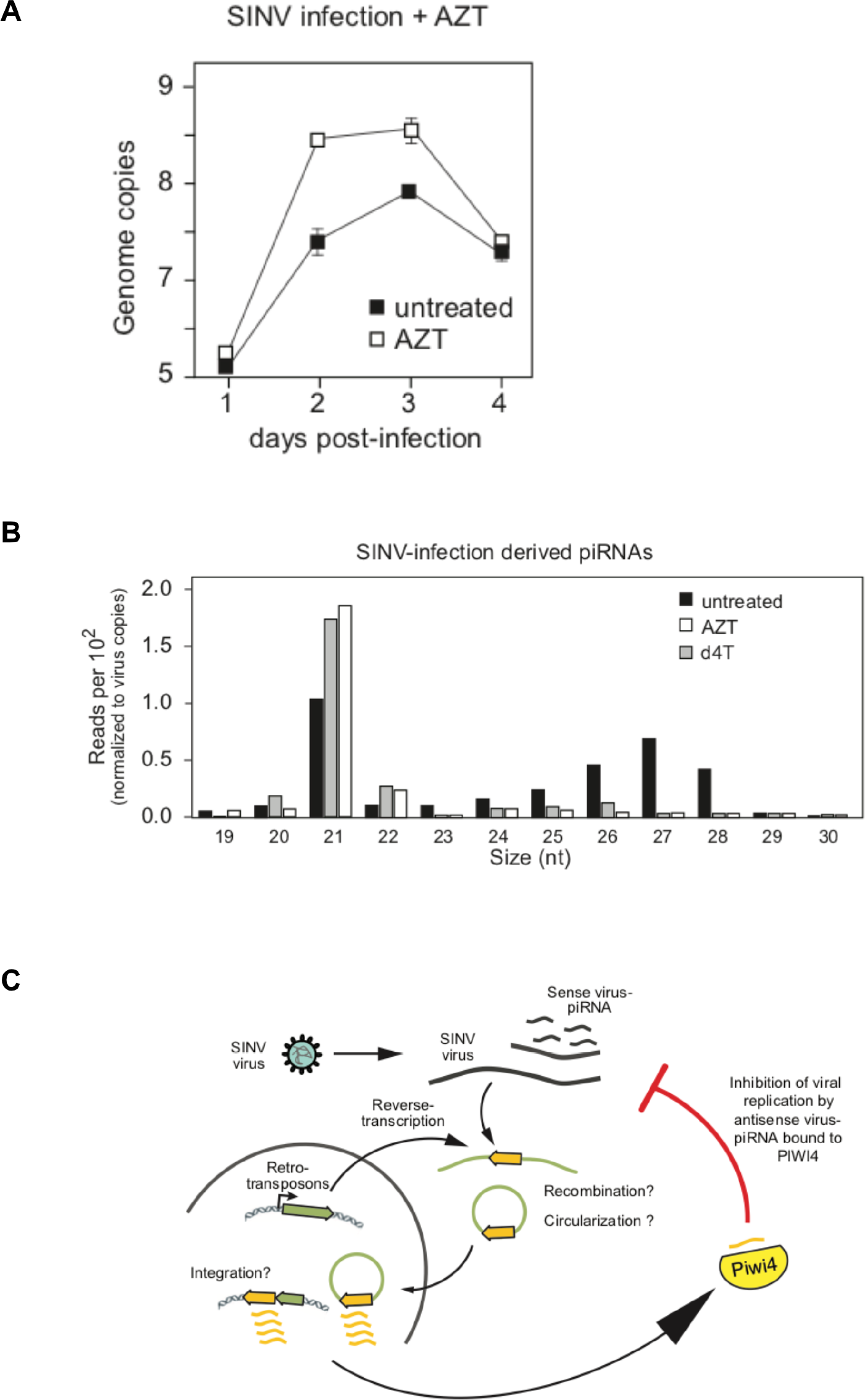

Recombination of a retrotransposon transcript with a foreign RNA molecule during the cytoplasmic stage of replication can occur during reverse-transcription by a copy-choice mechanism [30][31][32]. To determine whether recombination occurred between the RNA of an acutely-infecting arbovirus and a mosquito retrotransposon transcript, we analyzed the sequencing read pairs of circular DNAs to identify hybrids between virus and transposon sequences (Figure 5C). We found more than 200 hybrids between SINV and the most abundant LTR-retrotransposon sequences (Fig. 5E, 70 with Ty3 and 132 with Ty1 elements). Furthermore, mapping of hybrid reads from the most prevalent recombinants (SINV-Ty3/gypsy Element73) showed little recombination bias, nearly covering uniformly the entire SINV genome (Figure 5F, *i* and *ii*), which suggests that recombination is not driven by sequence homology. Unlike SINV, CFAV recombinants were found mostly associated with Ty1 elements (Figure 5 - supplement C). We also found direct evidence of uniform viral genome fragment conversion into circular vDNA (Figure 5 - supplement D *i* and *ii*) in DENV infected Aag2 cells.

Next, we examined whether vDNA synthesis and recombination with transposon elements can discriminate between self (host) and nonself (viral) RNAs. Only a small fraction of known expressed Aag2 mRNAs were found to have derived sequences in the episomal DNAs reads (~7.5% of known expressed mRNAs; 1093 out of 14612). Furthermore, there was no correlation between mRNA and episomal DNA abundance (R = 0.0045, Figure 5G and Figure 5 - supplement E). We detected no reads derived from the top 54 most abundant mRNAs, which account for 32% of all transcripts, and found no correlation between mRNA length and their enrichment in episomal DNA (R = 0.0786, Figure 5H). Indeed, the number of SINV vDNA reads was higher than for the most abundant host mRNA-derived episomal DNA (586 and 256 reads, for SINV and Aag2 transcript AAEL006357 respectively). Our results indicate that reverse transcription and recombination with retrotransposons is a highly specific mechanism, which preferentially converts viral RNA into vDNA. Together, these data suggest a model in which vDNA is acquired by the specific incorporation of viral RNA into retrotransposon replication complexes, followed by reverse transcription and recombination between the transposon and virus genome.

**Figure 5.**
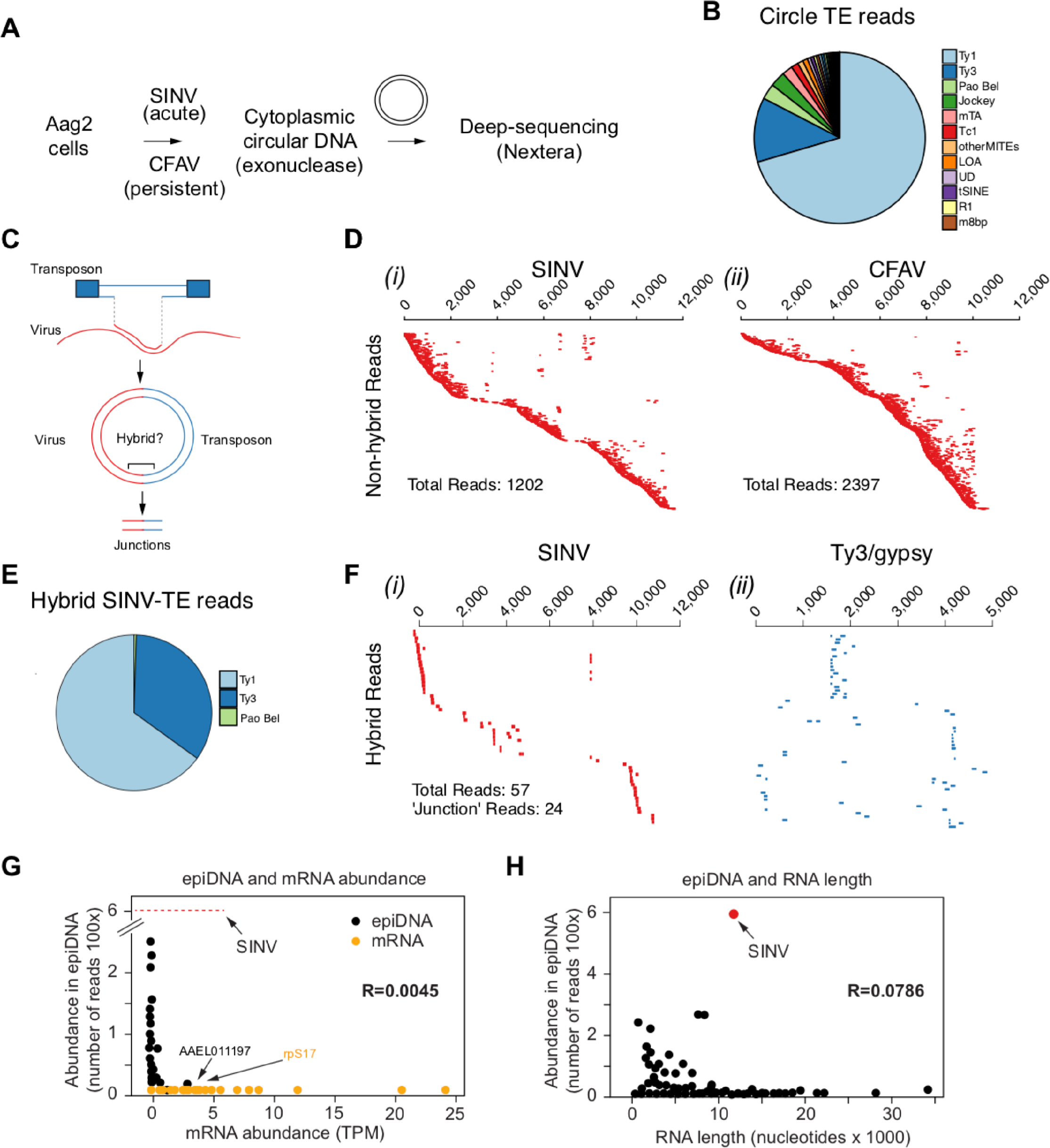

### Genomic analysis of Aag2 cells reveals independent acquisition of EVEs

If indeed reverse transcription and recombination are followed by integration into the host genome, this process could provide mosquitoes with a mechanism of immunological memory. Integration of vDNA derived from non-retroviral RNA viruses, known as endogenous viral elements (EVEs), have been identified in a wide variety of animal genomes[33][34][35] but the function, if any, of EVEs is still ill-understood. We considered that EVEs in the *Ae. aegypti* genome [34][36][37] may constitute a reservoir of virus sequences that can be utilized for the biogenesis of antiviral piRNAs. To examine this possibility, we sequenced the *Ae. aegypti* cell line Aag2 genome (~1.7Gb)[18]. The *Ae. aegypti* genome includes extreme heterozygosity, high repetitive content, and polymorphic inversions that together represent a significant challenge, particularly for the identification of newly acquired EVE sequences. We thus produced a high-quality assembly of the mosquito genome using single-molecule, real-time sequence technology (Pacific Biosciences sequencing) to generate long reads that permit the identification and analysis of highly repetitive regions (see Methods). Our data enabled an assembly that was over 10 fold more contiguous than the previous *Aedes* assembly[18], and permitted examination of highly repetitive regions. This was important because virus/retrotransposon hybrids are likely to insert into the genome at target sites to form repetitive transposon clusters[38]. Thus, analysis of our assembly led to the identification of new EVEs in the Aag2 genome.

Among the EVEs identified in our new Aag2 genome[18], we found a CFAV-derived EVE within contig 000933F. This fragment corresponds to CFAV NS2A gene coding region and it is flanked by long-terminal repeats (LTR) and *gag* Ty3-gypsy retrotransposon sequences (Figure 6A). This CFAV/Ty3 gypsy sequence is inserted into a piRNA producing cluster (piRNA cluster) among other LTR retrotransposons and EVEs. This piRNA cluster is transcriptionally active, producing antisense piRNAs targeting CFAV and members of other viral families, such as Rhabdoviridae and Reoviridae. Comparing this locus with the corresponding site in the *Ae. aegypti* Liverpool genome, from a strain isolated in West Africa, revealed that the CFAV EVE insertion observed in Aag2 cells is not present in the mosquito genome (Figure 6B), although other CFAV-derived EVEs are found at other loci in the Ae. aegypti AaegL3 assembly[17]. Of note, Aag2 cell line was originally established in 1968 in Israel from mosquito embryos[39]. Thus, these observations suggest that acquisition of EVEs is a dynamic process, perhaps depending upon the geographic location and infection history of mosquito populations[17] [40].

**Figure 6.**
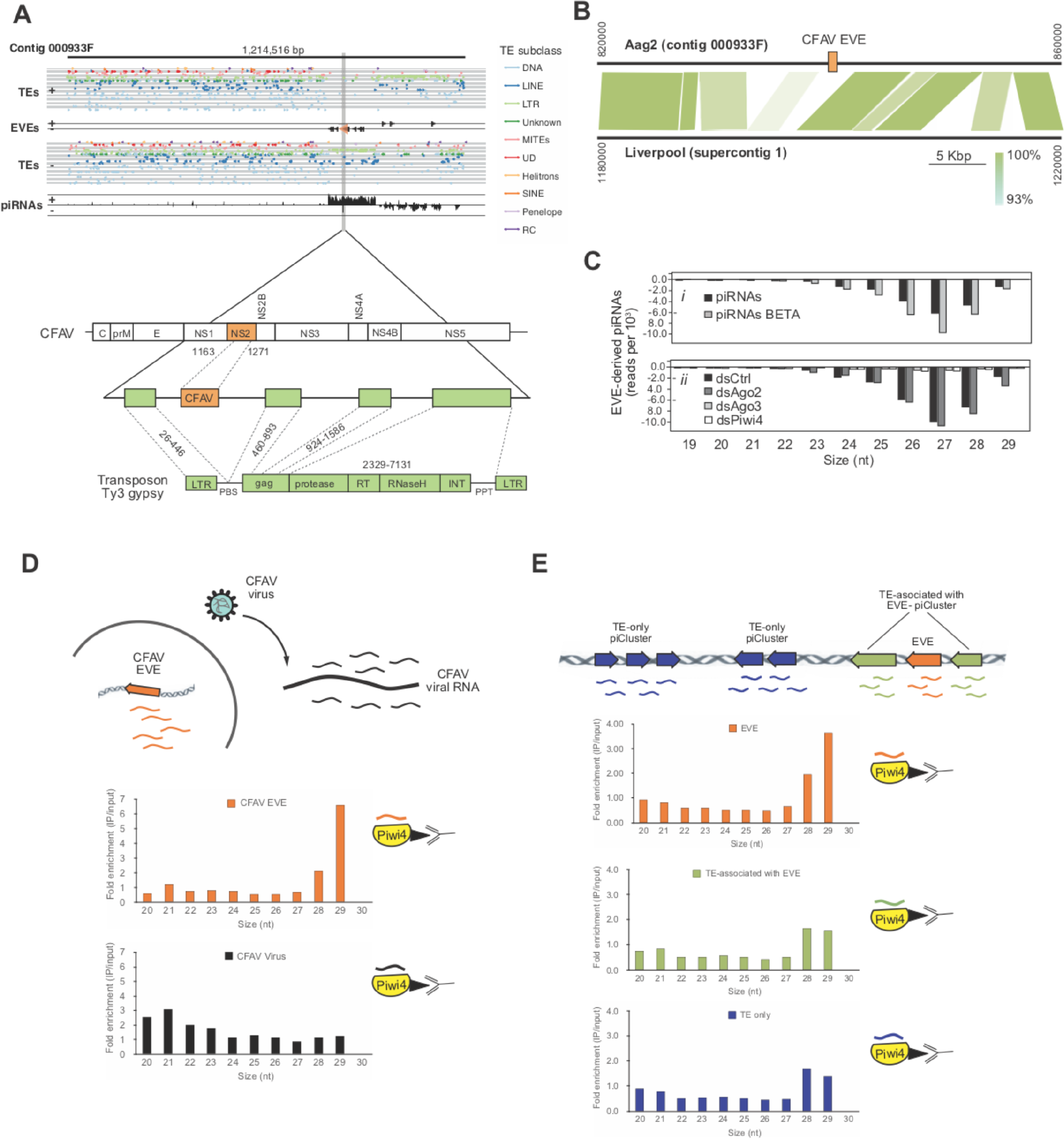

### Piwi4 binds specifically to antiviral piRNAs produced from EVEs

To test if EVEs are used to produce functional antiviral piRNAs, we analyzed EVE-derived small RNAs produced in Aag2 cells. The vast majority of small RNAs mapping to EVEs are 24-30 nt in length (>95%) and are almost exclusively antisense relative to the pseudo-ORF (>99%) (Fig. 6C, *i*). Furthermore, EVE-derived piRNAs showed a strong bias for uracil at the 1^st^ position (Figure 6 - supplement A) and were protected from beta-elimination, suggesting that they are mature piRNAs, methylated on their 3’end (Figure 6C, *i*). Importantly, knockdown of Piwi4 and Ago3, but not Ago2, resulted in a loss of EVE-derived piRNAs (Figure 6C, *ii*). Thus, the length, methylation, and sequence bias indicate that small RNAs derived from EVEs are *bona fide* piRNAs and their production require Piwi4 and Ago3. Because Piwi4 specifically binds to antisense v-piRNAs produced during acute infection (Figure 3A and C), we examined whether Piwi4 also preferentially associates with antisense piRNAs during persistent infection. Analysis of the new, highly contiguous, Aag2 genome assembly revealed that the existence of a unique CFAV-EVE in contig 933F, producing high level of antisense piRNAs (Figure 6A and B) and corresponds to the NS2a gene of CFAV strain distinct from the CFAV that established persistent infection in the Aag2 cell line (see Methods). Accordingly, the sequence differences between CFAV-EVEs and CFAV genomic RNA allowed us to distinguish between EVE- and virus-derived piRNAs. Strikingly, Piwi4-associated small RNAs mostly corresponded to CFAV EVE-derived antisense piRNAs and not to CFAV virus-derived piRNAs (Figure 6D and Figure 6 - supplement B) nor to TE-targeting piRNAs (Figure 6 - supplement C). Although most piRNAs are between 26 and 28 nt long, Piwi4 associated with significantly longer piRNAs (28-29 nt vs 26-27 nt, *X*^*2*^, *p* < 2.2e-16). Of note, during acute infection SINV-derived piRNAs bound to Piwi4 were also significantly enriched for longer sequences (Figure 3A). Again, the preferential binding of Piwi4 for longer piRNAs derived from genomic (EVE) versus RNA viral template was further confirmed in the other two replicates of the Piwi4-IP experiment (Figure 6 - supplement D). Similar results were obtained with the majority of piRNAs derived from EVEs identified in the Aag2 genome (Figure 6) and demonstrated that Piwi4 preferentially associated with longer EVE-piRNAs at the genome-wide level (28-29 nt vs 26-27 nt, *X*^*2*^, *p* < 2.2e-16).

The lack of enrichment in Piwi4 IP for the overall population of TE-targeting piRNAs (Figure 6 - supplement C) suggests that Piwi4 specifically discriminates between TE-targeting and EVE-derived piRNAs. We hypothesized that the signals allowing this specific piRNA sorting to Piwi4 might be contained in the genomic loci of EVEs. Because EVEs are strongly enriched in a sub-population of TE-rich piRNA clusters interspersed with LTR TE fragments[18], loci-specific sorting to Piwi4 may then extend to neighboring TE-targeting piRNAs (which would have been masked in the genome-wide analysis). We thus examined whether TE-targeting piRNAs transcribed from piRNA clusters associated with EVEs were preferentially bound to Piwi4 compared to TE-targeting piRNAs derived from piRNA clusters that contained only TE fragments. Like for piRNAs derived from TE-only loci, analysis of TE-targeting piRNAs transcribed from EVE containing piRNA clusters showed no enrichment for Piwi4 binding (Figure 6E). Therefore, piRNAs that associate with Piwi4 does not appear to be specified at the global piRNA cluster level. Taken together, these results demonstrate that Piwi4 binds preferentially to longer piRNAs and distinguishes between antiviral EVE-derived piRNAs and other piRNA cluster TE-targeting piRNAs through localized EVE-specific signals.

### EVE-derived piRNAs and Piwi4 provide antiviral defense against acute and persistent infections

Like antisense v-piRNAs from acute SINV infection in Aag2 cells, CFAV EVE-derived piRNAs are efficiently loaded onto Piwi4, which acts as a restriction factor against acute arboviral infections (Figure 1A and C). Analysis of sense/anti-sense CFAV piRNAs showed a characteristic ping-pong processing, 10-bp offset signature (Figure 7 - supplement A), suggesting that persistently infecting CFAV genome is cleaved by these anti-sense piRNAs (Figure 7 - supplement B). We thus sought to determine whether piRNAs derived from EVEs are capable of mediating silencing. To this end, we inserted into the 3’ UTR of a Renilla luciferase (Rluc) reporter gene ~500 bp sequences corresponding to *Ae. aegypti* genome-encoded EVEs in either the sense or antisense orientation (Figure 7A, *ii* inset). We selected four independent sequences with distinct piRNA expression levels (Figure 7A, *i* inset). Transfection of Rluc-EVE reporters into Aag2 cells demonstrated that Rluc expression was significantly reduced when the inserted EVE sequence was in the sense orientation as compared to the control antisense orientation (Figure 7A, *ii*). Furthermore, efficiency of silencing correlated with piRNA expression levels (Figure 7A, *i*). Thus, antisense EVE-derived piRNAs can silence mRNAs containing complementary sequences.

**Figure 7.**
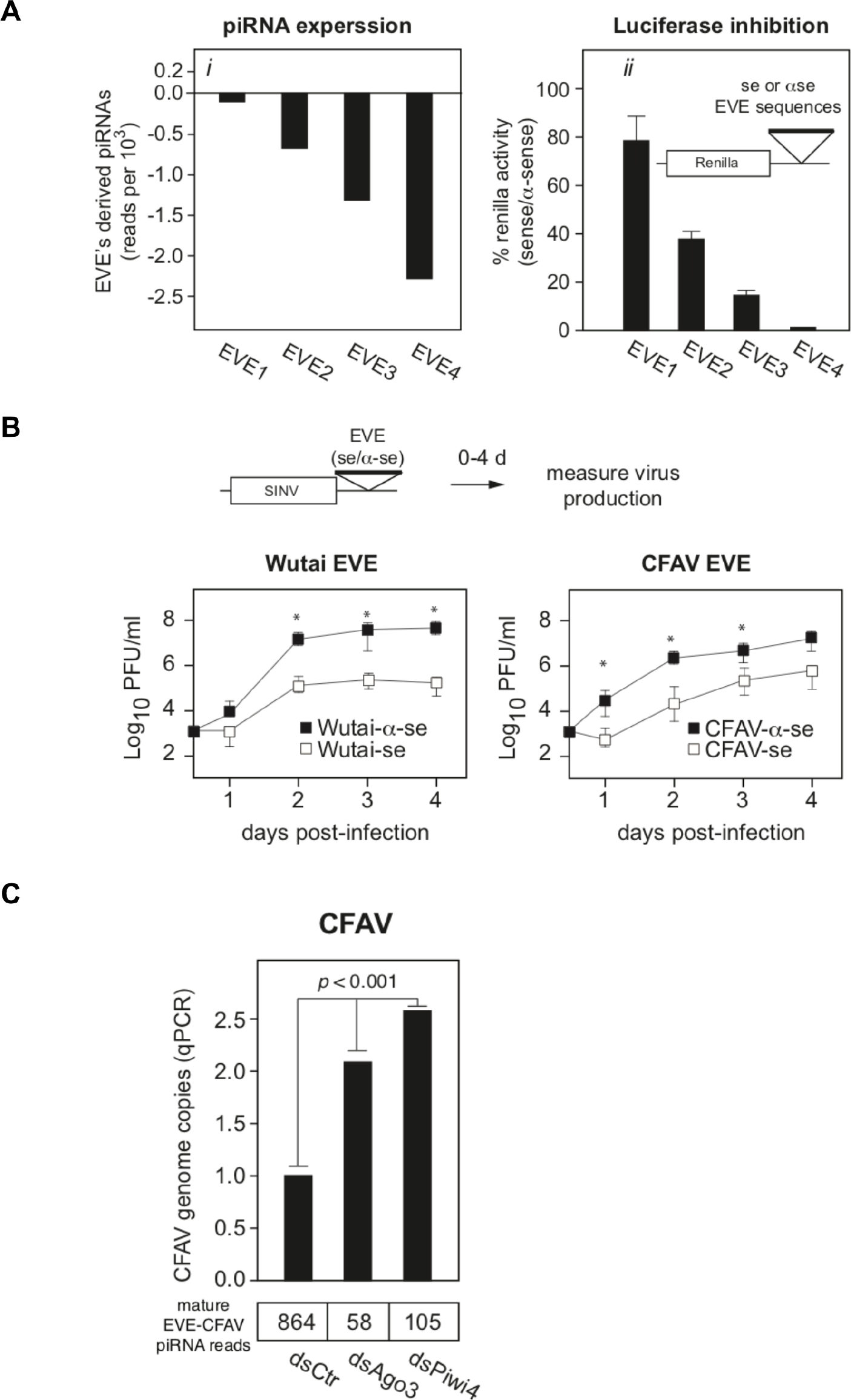

We next tested whether EVE-derived piRNAs can protect from virus infection. We first inserted two EVE sequences (derived from Wutai mosquito virus and CFAV) in the 3’UTR of SINV genome (Figure 7B). Infection of Aag2 cells with these viruses showed that insertion of EVE sequences in the sense orientation reduces virus replication 10 to 100 folds relative to the antisense EVE control viruses (Figure 7B, *p<0.*05 one-tail Wilcoxon rank sum test). We note that, while piRNAs that target the viral genome significantly decreased virus replication, it did not completely inhibit virus production. It is possible that the interaction of the EVE/piRNA system with the virus replication machinery may facilitate the establishment of a host-virus equilibrium typical of persistent infections. To test this hypothesis, we first determined the effect of Piwi4 knockdown on CFAV persistent infection and observed a significant increase in viral genome copy number (Figure 7C, *p*<0.001). Given that Piwi4 knockdown appears to affect both siRNA and piRNA production (Figure 2C and D), we sought to further examine the antiviral effect specific to piRNAs. To this end, we also performed Ago3 knockdown, which inhibit the production of piRNAs without significantly affecting siRNA levels (Figure 2C and 7C, bottom panel). Ago3 knockdown also increased CFAV genome copy number (Figure 7C, *p<0.*001). We thus conclude that mosquito EVEs generate piRNAs capable of effectively controlling virus replication.

## DISCUSSION

The immune system of *Aedes aegypti* mosquitoes allows arboviruses to establish persistent infection that facilitate spread in the human population. Investigating the role of the PIWI proteins in the mosquito immune response, we found that the anti-transposon surveillance pathway common to the majority of metazoan has evolved in mosquitoes as an effective adaptive antiviral defense. A central player on this pathway is Piwi4, which appears to specifically act in the mosquito antiviral piRNA system. In addition, episomal DNA and genome resequencing have revealed that conversion of viral RNA genomes into vDNA is a self-nonself discriminatory process and that EVEs-derived piRNAs are loaded onto Piwi4 to restrict acute and persistent infections in mosquito cells.

### The antiviral factor Piwi4 binds to v-piRNAs and is required for their maturation

RNAi is the main antiviral defense in insects. Unlike other diptera such as *Drosophila melanogaster*, arboviral infections of mosquito cells is known to lead to virus-derived piRNAs production[1], which are usually associated with genome surveillance against TEs[10]. In addition, expansion of the Piwi gene family in Aedes aegypti suggested functional diversification of its piRNA pathway. Indeed, studies of mosquito Piwi proteins have showed that Piwi5 and Ago3 bind to v-piRNAs and are involved in v-piRNA production but do not have antiviral activity. By contrast, Piwi4 was reported to control arboviral infections but not to bind to v-piRNAs nor contribute to their production. This discrepancy between antiviral function and piRNA production and binding has left our understanding of the piRNA machinery’s role in antiviral defense incomplete. Accounting for viral RNA level and piRNA methylation, we now demonstrate that in fact, Piwi4 binds to antisense v-piRNAs and is required for their production and maturation. In this context, the lack of antiviral effect in Piwi5 and Ago3 knock down experiments[8] suggests that v-piRNA accumulation from the ping pong amplification between Piwi5 and Ago3 might solely reflect a a degradation pathway for viral RNAs already silenced by other RNAi complexes, such as Piwi4 and Ago2.

In SINV-infected Aag2 cells, knock down of Ago2, which binds to v-siRNAs, led to a decrease of mature v-siRNAs but not of mature v-piRNA production. By contrast, knock down of Piwi4, which we found binds to v-piRNAs, led to a loss of both mature v-siRNAs and v-piRNAs. In addition, Piwi4 does not bind to TE-targeting piRNAs (Figure 3B) but is required for their maturation (Figure 2 - supplement F). Finally, our results and others[9] indicate that Piwi4 and Ago2 interact. Together, these data suggest that Piwi4 acts as a hub between the siRNA and piRNA antiviral pathways in mosquito and could affect siRNAs and piRNAs maturation by controlling the recruitment of the endogenous small RNA methyltransferase to RISCs. Ago3, which is also involved in TE targeting and v-piRNA methylation does not affect v-siRNA maturation and thus does not share the central role of Piwi4.

### Piwi4 binds to a specific form of v-piRNAs transcribed from vDNA elements

Production of vDNA molecules in *Aedes* mosquitoes during arboviral infection is linked to viral tolerance in mosquito vector[7] and consequently a capacity to act as an arbovirus vector. By sequencing episomal DNA in SINV and DENV infected Aag2 cells, we have confirmed that fragments of the infecting arbovirus are reverse-transcribed with LTR TEs providing unbiased coverage from loci across the entire viral RNA genome. Thus, recombination between viral genomes and transposons appears to be sequence-independent. Importantly, our analysis reveals that vDNA formation occurs through a self-nonself discriminatory process. An additional characteristic of a *bona fide* immune response to viral infection.

In the genome, vDNA sequences known as EVEs produce antisense v-piRNAs that can restrict acute and persistent infections (Figure 7). During processing, Piwi4 binds specifically to antisense v-piRNAs produced during acute SINV infection and EVE-derived piRNAs. In both cases, Piwi4 associates preferentially with long forms of v-piRNAs (28-29 nucleotides in length). Furthermore, inhibition of vDNA production leads to increased viral replication and decreased v-piRNA accumulation during acute SINV infection, similar to Piwi4 knock down. Because of this phenocopy between vDNA inhibition and Piwi4 knock down and the conserve binding preference of Piwi4 to v-piRNAs derived from EVEs and acute infection, we propose that antisense v-piRNAs bound to Piwi4 during acute arborviral infection derived from vDNA transcription rather than from the viral RNA itself. Consistent with this model, the lack of effect of Ago2 knock down on v-piRNA production despite Ago2 interaction with Piwi4, further suggests that v-piRNAs bound to Piwi4 are not processed from viral RNA genomes cleaved by Ago2-RISCs.

In Aag2 cells, EVE-containing piRNA clusters produce more piRNAs than loci without EVEs[18]. However, little is known about the regulatory mechanisms behind EVE-derived piRNAs production. Preferential binding of Piwi4 to EVE-derived v-piRNAs does not extend to neighboring TE-targeting piRNAs (Figure 6E). Therefore, sorting of EVE-piRNAs to Piwi4 might be regulated in a target-specific manner, that is able to discriminate between TE and virus sequences at the genomic level. In *Drosophila*, heterochromatin-dependent recruitment of the transcription machinery allows internal initiation within piRNA clusters and specific transcription of TE-targeting piRNA clusters versus TEs. Similarly, we speculate that epigenetic marks on EVEs might control their transcription and the specific sorting of EVE-derived piRNAs to Piwi4.

### Endogenized Viral Elements in the mosquito genome provide transgenerational antiviral immunity

Our study suggests that in *Aedes* mosquitoes, the piRNA pathway has acquired a novel antiviral function with specialized Piwi proteins, such as Piwi4, to restrict viral replication in somatic tissues of infected mosquitoes. Antiviral piRNA biogenesis is initiated by the formation of EVEs that is dependent on RT-mediated vDNA synthesis and integration into the genome (Figure 4B and 6B and C). EVEs derived from vertically transmitted insect-specific viruses (ISV) [18][17], are prevalent in the mosquito genome. Indeed, germline infection by ISVs could facilitate EVE endogenization and vertical transmission. Phylogenetic and phenotypic analysis of ISVs and arboviruses from the Bunyaviridae and Flaviviridae families suggests an arthropod origin to arboviruses[41]. In addition, ISV persistent infections in mosquitoes can restrict replication of some arboviruses[42]. Our results indicate that EVE-derived piRNAs decrease the replication of viruses with complementary sequences. In this context, we speculate that EVE-derived RNAi immunity against ISVs could force them away from the germline and drive a change from vertical to horizontal transmission leading to the emergence of arboviruses.

Conversely, for arboviruses that are already in circulation in vertebrate hosts and do not infect the mosquito germ line, additional mechanisms must exit to ensure maintenance of viral tolerance in the mosquito vector. Our work in adult female *Ae. aegpyti* shows that Piwi4 is upregulated in the midguts and carcasses after blood meal but not after sugar meal, suggesting that Piwi4 expression is stimulated in conditions associated with arbovirus exposure *in vivo*. Also, Piwi4 binds specifically to antisense v-piRNAs derived from vDNA elements (Figure 6D). In addition, inhibition of vDNA formation *in vivo* leads to *Ae. Aegypti* hypersensitivity to CHKV infection[7]. Together, these results suggest that exposure to circulating arboviruses could trigger an antiviral piRNA response in somatic tissues, consistent with the establishment of arbovirus persistent infection.

Finally, maternal inheritance of cytoplasmic piRNAs from active piRNA loci can convert homologous inactive clusters into strong piRNA producing loci in *Drosophila*[43]. Similar cross-talk between EVEs and newly acquired vDNA from infecting arboviruses with partial identity to EVEs could suffice to trigger piRNA production from vDNA and thus stimulate efficient antiviral activity through v-piRNAs and Piwi4. We propose that EVEs and Piwi4 are functional extensions of the piRNA pathway and provide a conserved mechanism of transgenerational adaptive antiviral immunity in mosquito.

## Materials and methods

### Cell culture and virus propagation

*Aedes aegypti* Aag2 cells were cultured at 28 °C without CO_2_ in Schneider’s *Drosophila* medium (GIBCO-Invitrogen), supplemented with 10% heat-inactivated fetal bovine serum (FBS), 1% non-essential amino acids (NEAA, UCSF Cell Culture Facility, 0.1 μm filtered, 10 mM each of Glycine, L-Alanine, L-Asparagine, L-Aspartic acid, L-Glutamic Acid, L-Proline, and L-Serine in de-ionized water), and 1% Penicillin-Streptomycin-Glutamine (Pen/Strep/G, 10,000 units/ml of penicillin, 10,000 μg/ml of streptomycin, and 29.2 mg/ml of L-glutamine, Gibco).

SINV (J02363) and CHIK (vaccine strain 181/25) stocks were produced by infecting Vero cells at low MOI (below 1) in MEM with 2% heat-inactivated FBS. After CPE was observed (~72 hours post infection), the supernatant was centrifuged at 3,000 g for 10 min, passed through a 0.45 μm filter, supplemented with 10% glycerol, flash frozen, and stored at −80 °C. Recombinant SINV-EVE strains were generated by inserting after an additional sub genomic promoter, a sequence cloned from a Wutai Mosquito virus-derived EVE (328bp) or CFAV-derived EVE (332bp) in the Aag2 cells genome. Cloning was performed using InFusion (Clontech) after PCR amplification of EVE regions in the sense (oligos IF5/IF6 for CFAV-EVE, IF9/IF10 for WutaiVirus-EVE) and antisense orientation (oligos IF7/IF8 for CFAV-EVE, IF11/IF12 for WutaiVirus-EVE) IF5_AagEVE_CFAV-NS2a_F ACCACCACCTCTAGATGGGTGTTGCTAGTGGCG IF6_AagEVE_CFAV-NS2a_R GGATCCATGGTCTAGTGCGGCCGCTCTTCCACCCCATTATCAGGC IF7_AagEVE_CFAV-NS2aAS_F ACCACCACCTCTAGATCTTCCACCCCATTATCAGGC IF8_AagEVE_CFAV-NS2aAS_R GGATCCATGGTCTAGTGCGGCCGCTGGGTGTTGCTAGTGGCG IF9_AagEVE_WutaiV_F ACCACCACCTCTAGACAATAATCTCAATGATGTCCTCGCG IF10_AagEVE_WutaiV_R GGATCCATGGTCTAGTGCGGCCGCCGCTATTGGA CAGATTGTAGACTGT IF11_AagEVE_WutaiV_AS_F ACCACCACCTCTAGACGCTATTGGACAGATTGTAGACTGT IF12_AagEVE_WutaiV_AS_R GGATCCATGGTCTAGTGCGGCCGCCAATAATCTCAATGATGTCCTCGCG DENV2 (Thailand 16681 for cell culture experiments and Jamaica 1409 for *in vivo* experiments) stocks for cell culture studies were produced as described above except that Huh7 cells were used for propagation and the viral stock was supplemented with 20% FBS instead of glycerol. DENV2 for mosquito studies was propagated in C6/36 (*Ae. albopictus*) cells cultured in Modified Eagle’s medium (MEM) supplemented with 7 % heat inactivated (56°C for 30 minutes) fetal bovine serum (FBS), 1% penicillin/streptomycin, 1% glutamine, 1% NEAA and maintained at 28°C at 5%CO_2_.

### Virus titration

Virus was titrated by plaque assay by infecting confluent monolayers of Vero cells with serial dilutions of virus. Cells were incubated under an agarose layer for 2 to 3 days at 37°C before being fixed in 2% formaldehyde and stained with crystal violet solution (0.2% crystal violet and 20% ethanol). DENV2 plaque assays were performed in LLC-MK2 cells in 24-well plates. Ten-fold serial dilutions of whole mosquito homogenate supernatant were added for 1 hour and cells were overlaid with agar. After 7 days of incubations at 37°C cells were stained by addition of 3mg/ml MTT (3-[4,5-dimethylthiazol-2-yl]-2,5-diphenyltetrazolium bromide) solution to the plate and incubated for 4 hours [44][45]. Viral titers were calculated, taking into account plaque numbers and the dilution factor.

### dsRNA preparation

PCR primers including the T7 RNA polymerase promoter were used to amplify *in vitro* templates for RNA synthesis using Phusion polymerase (NEB). Manufacturer’s recommendations were used for the concentrations of all reagents in the PCR. Primers were synthesized by *Integrated DNA Technologies*, Inc. (IDT). The thermocycling protocols are as follows: 98°C 2:00, (98°C 0:15, 65°C 0:15, 72°C 0:45, these three cycles were repeated 10X with a lowering of the annealing temperature by 1°C per cycle); (98°C 0:15, 60°C 0:15, 72°C 0:45, these three steps were repeated 30X), 72°C 2:00. RNA was synthesized in a 100 μl *in vitro* transcription (IVT) reaction containing 30 μl of PCR product, 20 μl 5X IVT buffer (400 mM HEPES, 120 mM MgCl2, 10 mM Spermidine, 200 mM DTT), 16 μl 25 mM rNTPs, and 1 unit of T7 RNA polymerase. The IVT reaction was incubated at 37°C for 3-6 hours and then 1 μl of DNase-I (NEB) was added and the reaction was further incubated at 37°C for 30 min. The RNA was purified by phenol-chloroform-isoamyl alcohol followed by isopropanol precipitation. RNA was quantified using a Nanodrop (Thermo Scientific) and analyzed by agarose gel electrophoresis to ensure integrity and correct size.

### dsRNA soaking

Prior to dsRNA soaking, Aag2 cells were washed once with phosphate buffered saline w/o calcium or magnesium (dPBS, 0.1 μM filtered, 0.2g/L KH_2_PO_4_, 2.16g/L Na_2_HPO_4_, 0.2g/L KCl, 8.0g/L NaCl). Cells were soaked in 5 μg /ml dsRNA in minimal medium (Schneider’s *Drosophila* medium, 0.5% FBS, 1% NEAA, and 1% Pen/Strep/G) for the time indicated by the experiment. All incubations were performed at 28 °C without CO_2_. 3 days later, dsRNA-treated Aag2 cells were infected with SINV (MOI=1) and collected at 3 day post infection.

### Ae. aegypti

Adult *Ae. aegypti* (Chetumal strain) female mosquitoes were taken 2-3 days after emerging from the pupa stage and intrathoracically (IT) injected with 500 ng of dsRNA 2-4 days post-emergence. No randomization or blinding of the group allocation of mosquitoes was used for these studies. Three days later mosquitoes were infected with DENV-2 Jam1409 using an artificial bloodmeal consisting of DENV2-infected C6/36 (A. albopictus) cells in L15 medium suspension (60% vol), defibrinated sheep blood (40% vol); Colorado Serum Co., Boulder, CO) and 1 mM ATP. DENV2 titers in the bloodmeal were 1.0 ± 0.4 × 10^6^ PFU/mL. Blood fed females were selected, provided water/sugar and maintained in the insectary at 28oC, 82% relative humidity. For all mosquito experiments sample size was selected based on known variability from previous experiments.

Ovaries, midguts and carcasses were dissected at 0, 4, 7, 10 and 14 dpi and collected in 100μl of Trizol reagent (Invitrogen) and total RNA was extracted following the manufacturer’s instructions. Whole mosquitoes were also collected at 0, 4, 7, 10 and 14 dpi and triturated in 1 ml of supplemented MEM. After centrifugation the supernatant fluid was filtered (Acrodisc Syringe filters with 0.2 μm HT Tuffryn membrane) and virus titers were determined by plaque assay.

### Identification of CFAV strains

Primers (CFAV-fw and CFAV-rev) were designed to anneal to regions flanking the NS2A coding sequence of CFAV that are conserved between the Bristol (Gen-Bank# KU936054) and the Culebra (GenBank# AH015271.2) strains (Region 3321-3851 in Bristol coordinates). Sanger sequencing of the amplicon produced with these primers from cDNA generated from Aag2 cells identified the persistent infecting CFAV virus as belonging to the Bristol strain.

CFAV-fw: GCGAGGAACCAGAACCAACA

CFAV-rev: GCAGGACGCTCTTGTAGGC

### Luciferase assays

Cells were soaked in dsRNA for the indicated period of time using the dsRNA soaking method above. Prior to transfection, cells were washed 3X with dPBS, and then added to complete medium. Cells were transfected with plasmids encoding Firefly (pAc Fluc) and Renilla luciferase (pUb Rluc) with Effectene (Qiagen) using the manufacturer’s instructions with the following modification: 200 ng pAc Fluc and 50 ng pUb Rluc were used per transfection with a ratio of 1 μl effectene / 250 ng plasmid DNA. Firefly and *Renilla luciferase* sequences from the plasmids pGL3 and pRL-CMV (Promega) were cloned into pAc/V5-HisB (Invitrogen) and pUb (Ubiquitin promoter [46] respectively.

Twenty-four hours post transfection, cells were lysed in 50 μl passive lysis buffer (Promega), and Firefly and Renilla luciferase activity was determined from 10 μl of lysate using a dual luciferase reporter assay system using the manufacturer’s instructions (Promega) and analyzed on an Ultra-evolution plate reader (Tecan) using an integration time of 100 ms.

The analysis of effects of dsRNA uptake on target gene candidates was performed as above with the following exceptions: 30,000 cells were seeded per well, cells were treated with the initial dsRNA (targeting the candidate gene) for 72 hours, washed 3X with dPBS, before the addition of the secondary dsRNA (targeting the reporter).

### qPCR Analysis

Total RNA was extracted using TRIzol (Life Technologies). cDNA synthesis was performed using the iScript cDNA synthesis kit (Bio-Rad). Primers for RT-qPCR were obtained from IDT and are listed in the supplementary material. Specific genes or viral genomes were analyzed using SYBR green methods on a CFX- Connect (Bio-Rad). All genes/viruses tested for relative quantitation were normalized to RP49 expression. Relative quantitation was calculated by the 2^-ddCt^ method. Absolute quantitation was calculated using a standard of known quantity.

### Statistical analysis

Values were expressed as means +/− standard deviation. RNA expression levels (viral genome copies and endogenous mRNA) among samples were not assumed to have normal distribution and therefore were analyzed using the non-parametric test a two-tailed Mann-Whitney U-test to test for the likelihood of the RNA level in the treated condition is different from the control.

Comparison of growth curves of Sindbis virus strains containing a sense versus antisense cognate sequence to endogenous EVE-derived piRNAs were performed on the mean values of three independent infection experiments using the non-parametric (no assumption of normal distribution) Mann-Whitney-Wilcoxon test. Viral strains with sense targets for antisense EVE-piRNAs were expected to have decreased titers and therefore one tailed tests were used.

Chi square (*X*^*2*^) tests were performed on 2×2 contingency tables for the different size or strandness of v-piRNAs using the chisq.test in R.

### Affinity purification (AP)

5 × 10^6^ Aag2 cells were seeded in 10 cm dishes and allowed to attach overnight. Cells were transfected with expression plasmids using Transit2020 (Mirius Bio) using the manufacturer’s instructions. 24 hours post transfection cells were washed with dPBS three times, scraped off the dish in IP- buffer (pH 7.5 @ 4°C, 10 mM Tris, 2.5 mM EDTA, 250 mM NaCl, and complete protease inhibitor, Roche), and centrifuged at 2000 rcf for 5 min at 4°C. Cell pellets were resuspended in 300 μl lysis buffer (IP-buffer + 0.5% NP-40) and incubated at 4°C for 30 min and then centrifuged at @ 12000 rcf for 10 min at 4°C. The supernatant was added to 50 μl of Protein A conjugated beads (Sigma) and rotated for 1 hr at 4°C. Lysate was adjusted to 1% NP40 (by adding 4 volumes IP-buffer) and then transferred to 50 μl of anti-FLAG conjugated beads (Sigma, Cat #F2426) and rotated for 6-16 hr at 4°C. Beads were then washed 6 times with wash-buffer (IP-buffer + 0.05% NP- 40). Following the final wash 30 μl of elution buffer was added (IP-buffer + 100 μg/ml 3x Flag peptide, Sigma) and rotated for 1 hr at 4°C.

### Mass Spectrometry

To ensure samples were appropriate for mass spectrometry, eluates from affinity purification were analyzed by western blot and silver stain (Pierce). Samples were run on an Orbitrap LC-MS and analyzed with MaxQuant software.

### Piwi4 associated piRNAs

Piwi4 Flag was overexpressed in Aag2 cell and immuni-precipitated from SINV infected Aag2 cells at 3 dpi. After three washes with TBS + 1mM EDTA and 80 U/mL murine RNase Inhibitors (NEB), 1 ml of TRIzol was added to Flag beads and RNA extraction was performed according to the manufacturer’s protocol. Cloning of small RNAs was performed as described below.

The additional Piwi4 pull down experiments were performed similarly except for the following. After the three washes, small RNAs were first released from the beads by adding 20 mg/mL Proteinase Kand 80 U/mL murine RNase Inhibitors for 1h at 55°C, followed by TRIzol extraction. Cloning of the small RNAs were performed as described below except that there was no size selection for the bound fraction and ligation of the 3’ adapter was performed overnight at 18°C for increase efficiency.

For the eGFP control pull down experiment, a pUb-eGFP plasmid was transfected instead of the pUb-Piwi4:FLAG. Pull down and small RNA extraction and cloning were performed as described for the additional Piwi4 pull down experiments.

### Small RNA cloning for deep sequencing

7 × 10^6^ Aag2 cells were seeded in each T-75 flask in complete medium and allowed to attach overnight. Cells were washed with dPBS three times, scraped off the dish in dPBS, and centrifuged at 2000 rcf for 5 min at 4°C. RNAs were isolated using the miRvana kit (Life technologies). The large RNA fraction was used for RT-qPCR. The small RNA fraction was precipitated by adding 1/10^th^ volume 3M NaOaC pH 3.0, 1 μL gylcoblue (Life technologies), and 2.5 volumes 100% EtOH and incubated at −80°C at least 4 hours and then centrifuged at 12000 rcf for 10 min at 4°C. The pellet was washed with 80% EtOH and then resuspended in Gel Loading Buffer II (Life Technologies) and run on a 16% polyacrylamide gel containing 8M urea. Small RNAs (17-30 nt) were cut out from the gel and eluted overnight at 4°C and precipitated by adding 1/10^th^ volume 3M NaOaC pH 3.0, 1 μL gylcoblue (Life technologies), and 2.5 volumes 100% EtOH and incubated at −80°C at least 4 hours and then centrifuged at 12000 rcf for 10 min at 4°C. Small RNAs were cloned using microRNA Cloning Linker1 (IDT) for the 3’ ligation (ligation at 25°C for 3h) and the modified 5’ adapter (ligation at 37°C for 2h) with randomized 3’-end (CCTTGrGrCrArCrCrCrGrArGrArArTrTrCrCrArNrNrNrNrN). Libraries were run on a HiSeq 1500 or 2500 using the Rapid run protocol.

For circular DNA deep-sequencing, cytoplasmic fractions were isolated from Aag2 cells and processed for DNA extraction using a Nucleospin tissue kit (Macherey-Nigel). DNA was treated overnight with Plasmid-Safe ATP-dependent-DNase (Epicentre) overnight at 37 ̊C to remove non-circular DNA. 1ng of Plasmid-Safe treated DNA was used for Nextera cloning (Nextera DNA Library sample prep kit, Illumina) and sequenced on an HiSeq 4000. Bioinformatic analyses were carried out as described[47]. More than 50% comes from the genome (but are not mitochondrial nor TE derived). Of that 50%, about 1% comes from transcribed regions, suggesting very little contamination from cellular transcripts.

### Small RNA bioinformatics

Adaptors were trimmed using Cutadapt[48] with the --discard-untrimmed and -m 19 flags to discard reads without adaptors and below 19 nt in length. Reads were mapped using bowtie[49] using the −v 1 flag. Read distance overlaps were generated by viROME[50]. Sequence biases were determined by Weblogo[51]. Transposon sequences for *Ae. aegypti* were downloaded from TEfam (http://tefam.bio-chem.vt.edu/tefam/get_fasta.php) Sequencing depth, virus and TE read counts are presented in the Table 1 and 2.

**Table 1.**

| normal | dsCtr | dsAgo2 | dsAgo3 | dsPiwi4 |
| --- | --- | --- | --- | --- |
| Total reads | 14779383 | 13697808 | 10207667 | 12139804 |
| virus | 194682 | 659432 | 125859 | 301905 |
| TE | 1617084 | 1220933 | 782789 | 1484997 |

**Table 2.**
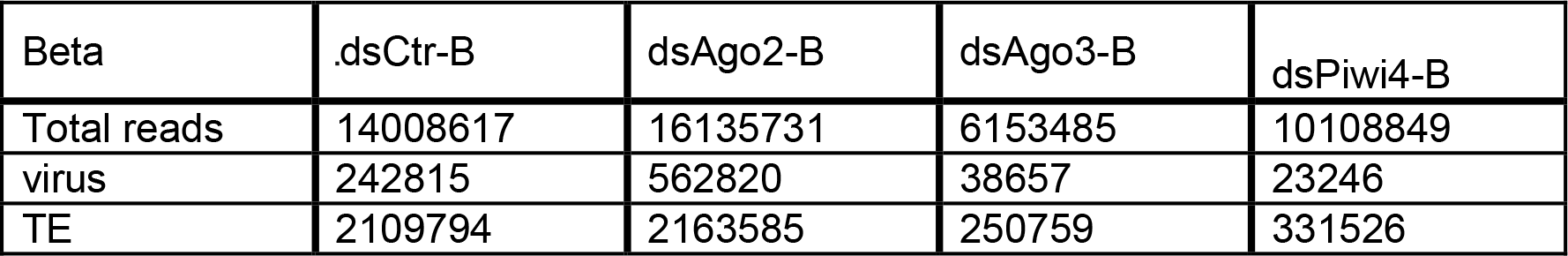

For the Piwi4 pull-down experiments, small RNA reads were mapped using Bowtie 1.2.1.1 in v mode, allowing up to 2 mismatches (-v 2). SINV and transposon-derived reads were mapped using the SINV genome (accession # NC_001547) and the TEfam database respectively. To specifically identify small RNAs derived from CFAV-EVEs, reads were first filtered out by mapping them to the persistently infecting CFAV virus genome (Bristol strain, accession # KU936054). Unmapped reads were then aligned to CFAV-EVEs identified in the PacBio Aag2 genome. Conversely CFAV-virus specific reads were identified by first filtering out CFAV-EVEs mapping reads, followed by the alignment of unmapped reads to CFAV virus genome (Bristol strain, accession # KU936054). For the genome-wide analysis of EVE-derived small RNAs, reads were aligned to all EVEs identified in the Aag2 PacBio genome without the CFAV-EVEs to avoid confounding effect from the previous analysis. For the three additional pull-down followed by small RNA-seq experiments, elimination of potential concatenate reads was achieved by only retaining reads that align perfectly (-v 0) to the different reference genomes of interest (e.g.: SINV, TEfam database) over their entire length, starting with a read length of 45 nucleotides. Reads that failed to align were trimmed of 1 base and assessed again for perfect alignment to the reference genome of interest. This process was repeated to the lowest read size of 18 nucleotides.

Reverse transcriptase inhibitor treatments was as carried out as described in[23].

### Transcriptome analysis of Aag2 cells

Aag2 transcripts abundance was calculated using published RNA-seq data from Aag2 cells (https://doi.org/10.1186/s12864-016-3432-5) and the Galaxy platform (usegalaxy.org, doi:10.1093/nar/gkw343). RNA-seq reads were aligned and quantified to the Liverpool *Aedes aegypti* transcripts reference (Aedes-aegypti-Liver-pool_TRANSCRIPTS_AaegL3.4.fa, available at www.vector_base.org) using the transcript quantification tool, Salmon[52]. Relative transcript abundance is expressed as Transcript Per Million. Comparative analysis of SINV and Aag2 mRNA-derived sequences in episomal/circular DNA sequencing dataset (see above) was performed using Bowtie 1.2.1.1. To filter out contamination from genomic sequences, episomal DNA reads were first aligned to the new Aag2 genome obtained by PacBio sequencing (see below), in v mode allowing up to 2 mismatches (-v 2). SINV and host mRNA-derived sequences were then identified by aligning the genome-unmapped reads to SINV genome (accession # NC_001547) or *Aedes aegypti* transcripts (Aedes-aegypti-Liverpool_TRANSCRIPTS_AaegL3.4.fa) respectively, using the same v mode (-v 2).

### EVE Identification

Identification of EVEs was achieved using standalone Blast. Blast Searches were run using the Blastx command specifying the genome as the query and a refseq library composed of the ssRNA and dsRNA viral protein-coding sequences from the NCBI genomes website as the database. The E-value threshold was set at 10^−6^.

The EVE with the lower E-value was chosen for further analysis to predict EVEs that overlapped. Several Blast hits to viral protein genes were identified as artifacts because of their homology to eukaryotic genes (e.g. closteroviruses encode an Hsp70 homologue). These artifacts were filtered by hand.

To determine the necessity of genes for the biogenesis of virus RNA-derived piRNAs, we knocked down expression of each gene in Aag2 cells by dsRNA soaking and infected with SINV (MOI=10). As a control, we soaked Aag2 cells in Fluc dsRNA. Following a four-day infection, we harvested the small RNA (<200 nt) for deep sequencing analysis and large RNA (>200 nt) for RT-qPCR analysis of knockdown efficiency and SINV genome copy number. RT-qPCR analysis of these samples confirmed that knockdown of each gene was effective (>90%) and SINV genome copy number increased with each treatment relative to control.

### PCR and Cloning

Genomic DNA was purified from Aag2 cells using Nucleospin Tissue mini spin columns (Macherey-Nagel). Fragments of EVEs were amplified from genomic DNA by PCR using Phusion polymerase (NEB). ~500 bp PCR products from EVEs were non-directionally cloned into the 3’ UTR of pUb-Renilla using NotI (NEB). Clones of inserts in both sense and antisense polarity were isolated and amplified.

## Data availability

Next generation sequencing libraries of small RNAs and extrachromosomal circular DNAs are available through NCBI Sequence Read Archive (SRA) at https://www.ncbi.nlm.nih.gov/sra/PRJNA493127.

---

**Figure 1 - Supplement.**
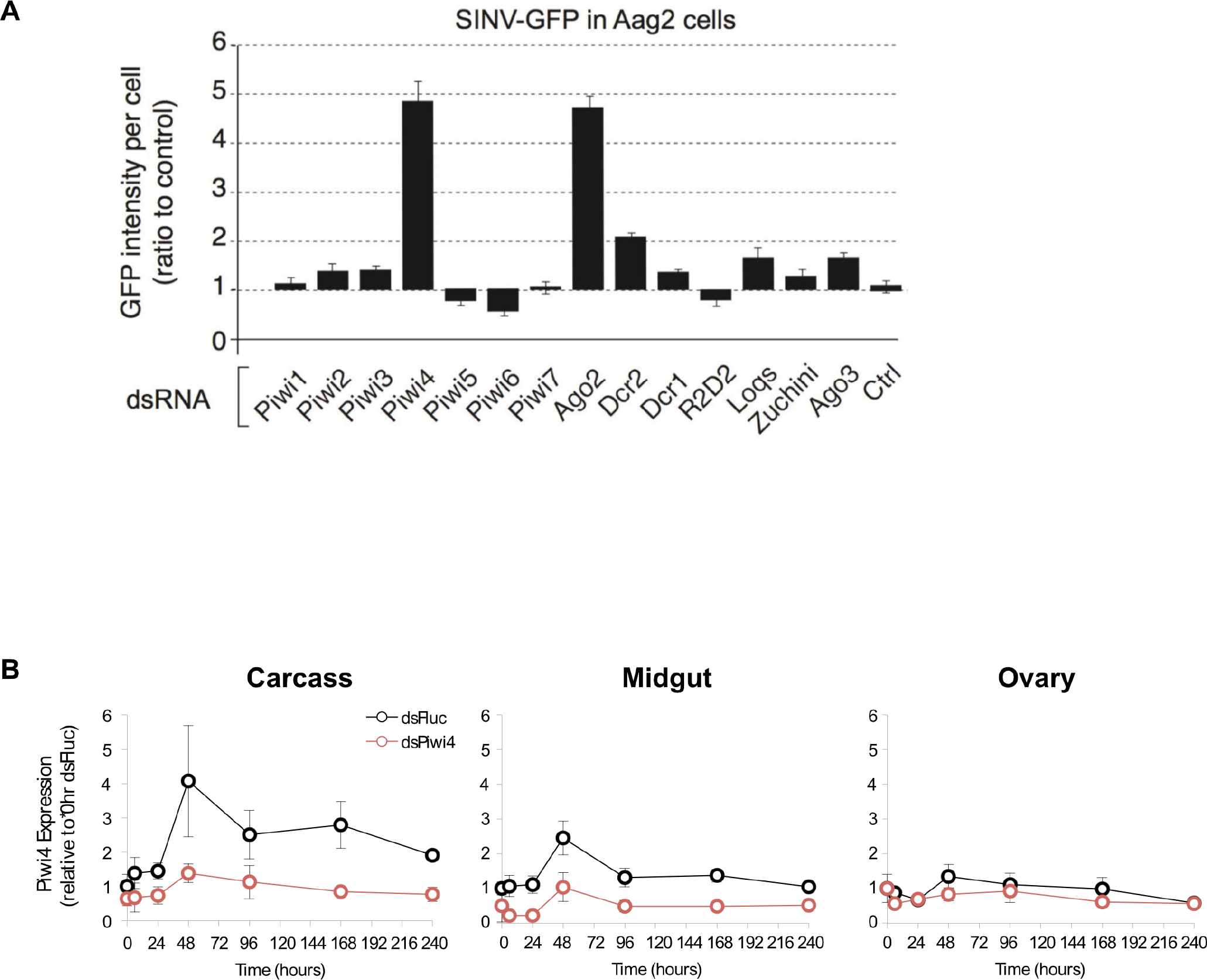
Piwi4 knockdown in cell culture and adult female *Ae. aegypti* mosquitoes. **A.** RNAi screen against the seven *Aedes aegypti* Piwi genes and other selected RNAi factors in SINV-eGFP infected Aag2 cells. Viral replication was measured as GFP intensity per cell. **B.** Piwi4 mRNA measured by qPCR from pools of dissected tissues from *A. aegypti* mosquitos injected with dsRNA against Firefly luciferase (dsFluc, negative control) or against Piwi4 (dsPiwi4). The error bars represent standard deviations of 4 biological replicates of pools of 5 of the respective tissue. Significant changes over control are marked with asterisks (p ≤ 0.05, Mann-Whitney U test).

**Figure 2 - Supplement.**
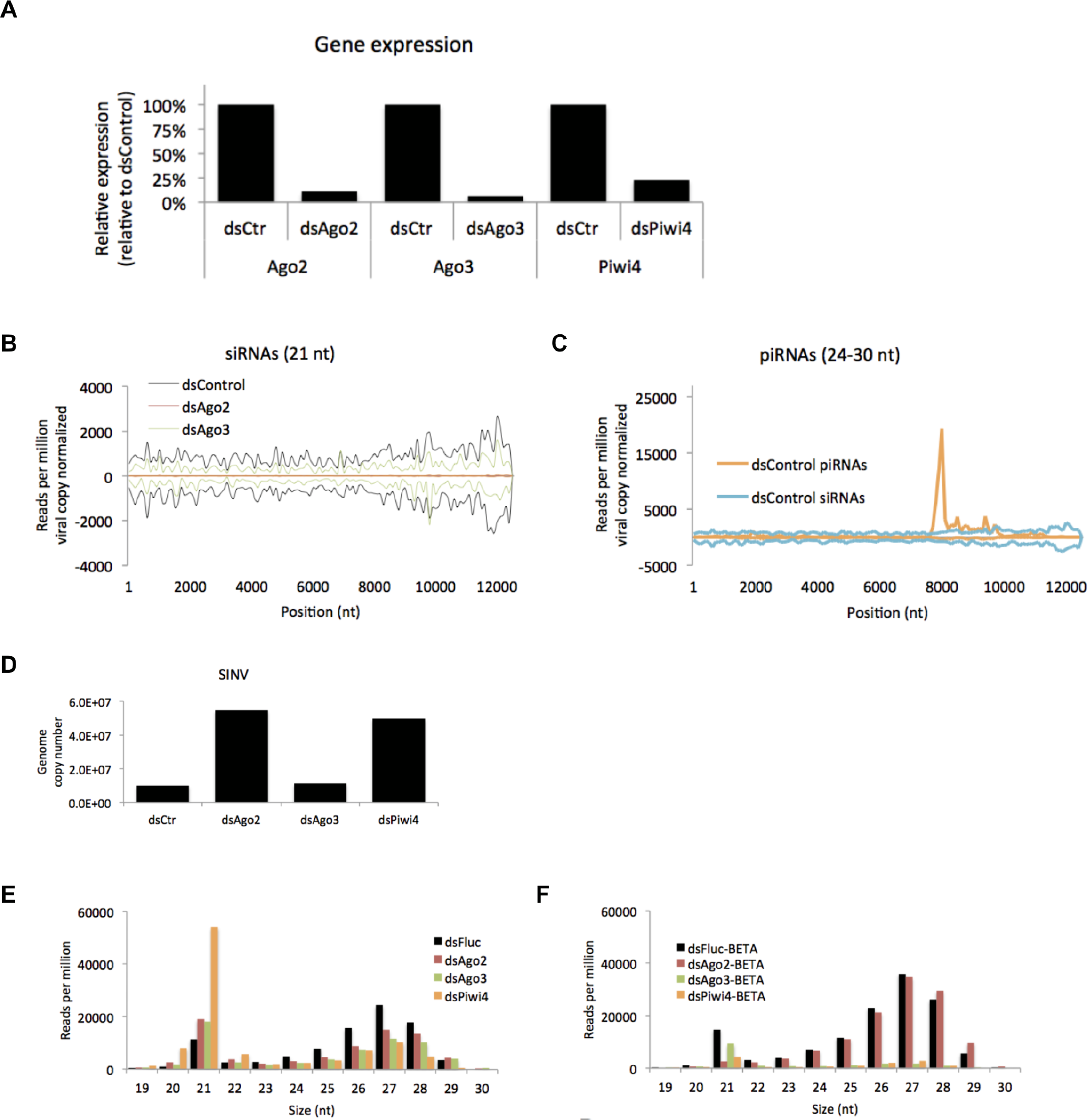
Effects of Ago2, Ago3 and Piwi4 knockdown on SINV-derived small RNAs. **A.** Ago2, Ago3, and Piwi4 mRNA measured by RT-qPCR from large RNA fraction of sample used for small RNA library preparation. **B.** Mapping of siRNAs (21 nt) to the SINV genome from infected Aag2 cells treated with control, Ago2, Ago3, or Piwi4 dsRNA. **C.** Mapping of piRNA (24-30 nt) to the SINV genome from infected Aag2 cell treated with control dsRNA. **D.** SINV genomic RNA measured by RT-qPCR from large RNA fraction of sample used for small RNA library preparation. **E.** Size distribution plot of small RNA mapping to transposons from infected Aag2 cell treated with control, Ago2, Ago3, or Piwi4 dsRNA. **F.** Size distribution plot of beta-elimination resistant small RNA mapping to transposon from infected Aag2 cells treated with control, Piwi4, or Ago3 dsRNA.

**Figure 3 - Supplement.**
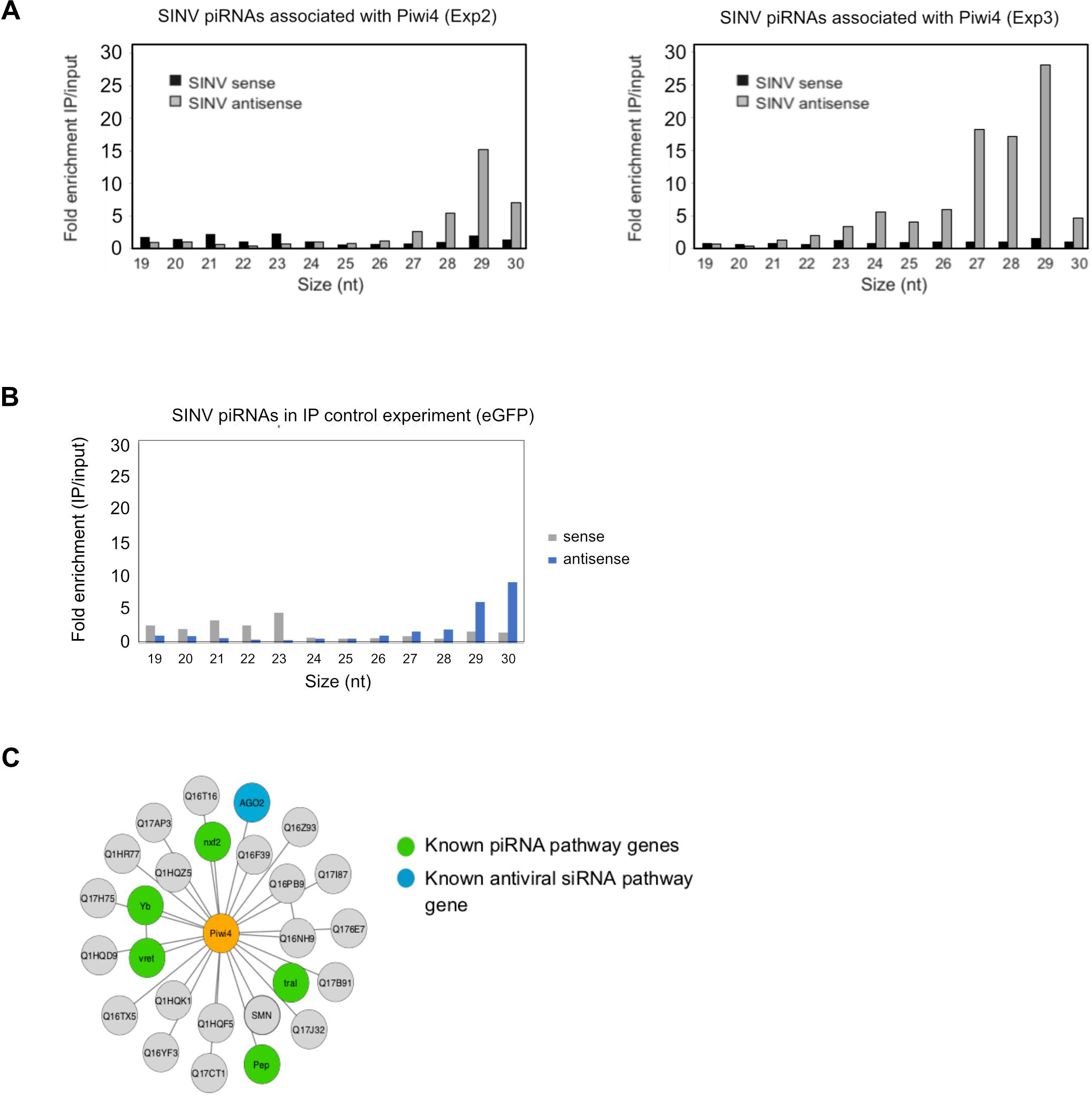
Piwi4 binds to bona-fide v-piRNAs and interacts with known siRNA and piRNA pathway components. **A.** Results from two independent biological replicates (Exp1 and Exp2, left and right panel respectively) for the enrichment of SINV-derived small RNA in Piwi4-IP fraction compared to input sample. **B.** Lack of enrichment of SINV-derived small RNA in control eGFP immunoprecipitation (IP) fraction compared to input sample. **C.** Network of Piwi4 protein interactions identified by affinity purification followed by mass-spectrometry using a C-terminal FLAG tagged Piwi4 expressed in Aag2 cells as bait. Proteins known to be involved in piRNA pathways or in antiviral response in insects are shown in green or blue respectively.

**Figure 4 - Supplement.**
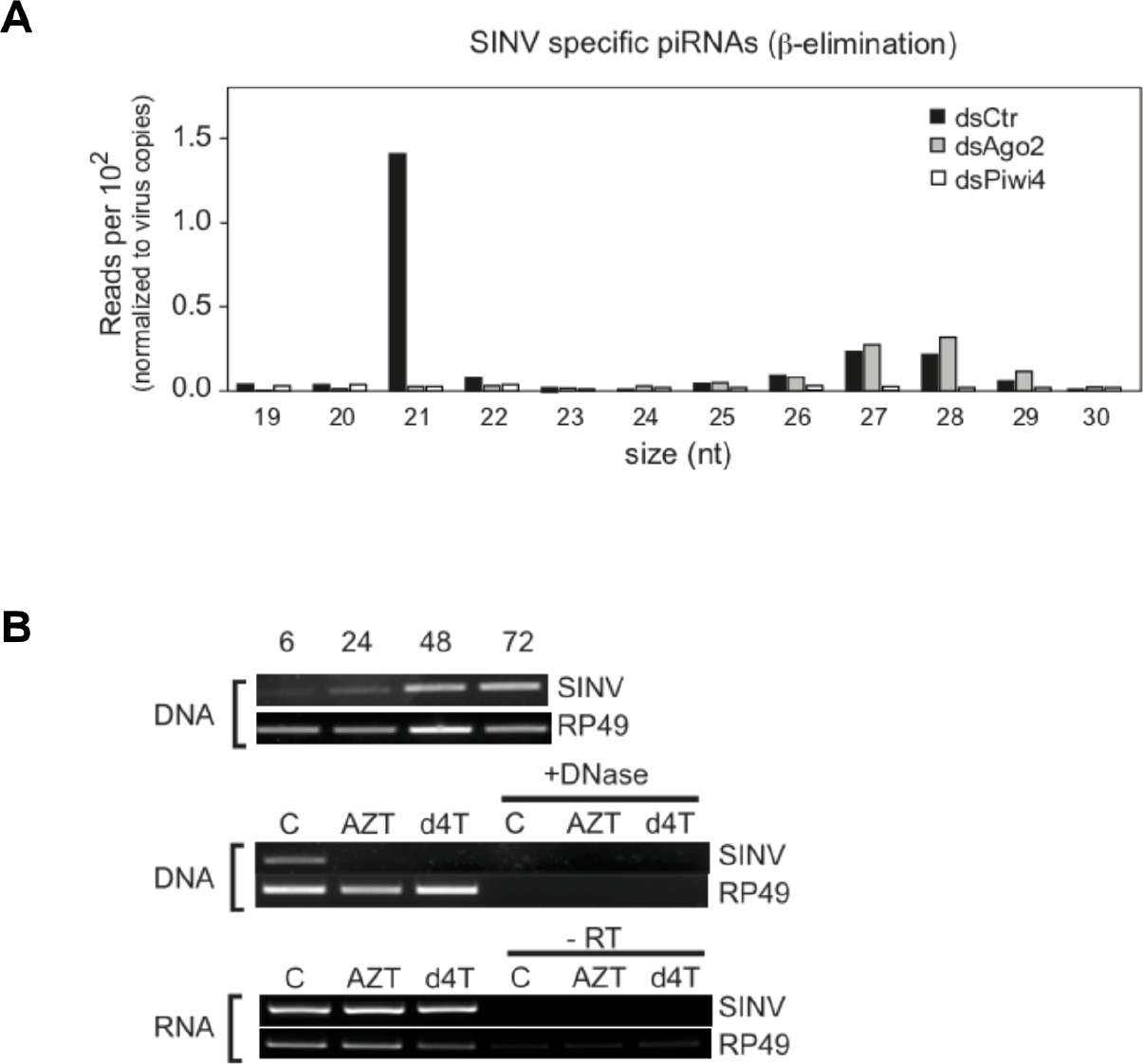
Piwi4 binds to bona-fide v-piRNAs and interacts with known siRNA and piRNA pathway components. **A.** Size distribution plot of ß-elimination resistant small RNAs mapping to the SINV genome from infected Aag2 cells treated with control, Piwi4 or Ago2 dsRNA. Read counts per hundred were normalized to SINV genome copy numbers. **B.** Detection of SINV vDNA (PCR) and viral RNA (RT-PCR) in extracted RNA or genomic DNA from infected Aag2 cells over time or in infected Aag2 cells treated with either AZT or d4T. DNase I treatment prior to PCR prevents vDNA detection. Reactions lacking reverse transcriptase were used as a negative control for viral RNA detection.

**Figure 5 - Supplement.**
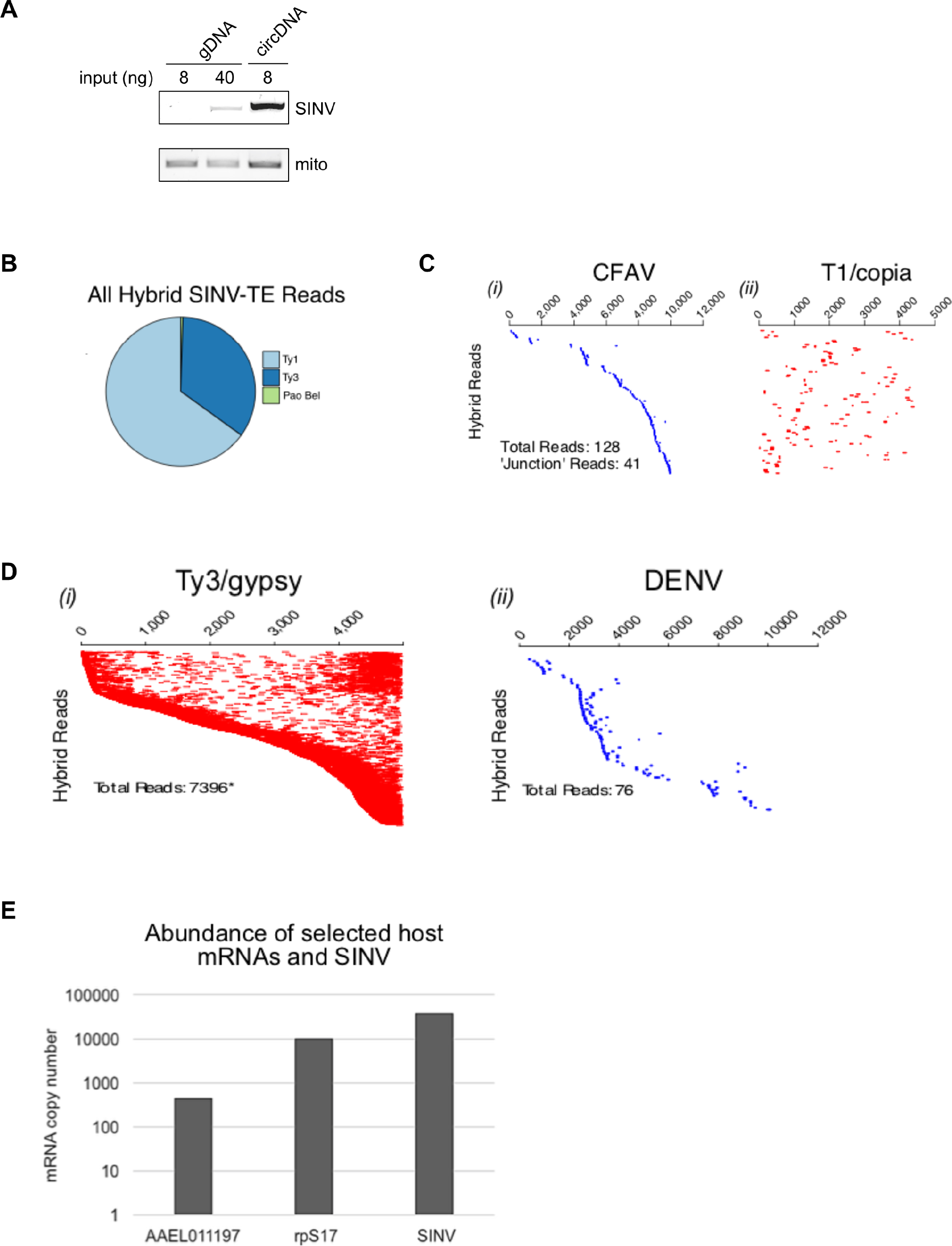
Supplement. Analysis of episomal viral and transposon-derived DNA in arbovirus-infected Aag2 cells. **A.** SINV infected-Aag2 cells were collected at 4 d.p.i and processed for genomic DNA (gDNA) or episomal circular DNA (circDNA) extraction. 8 or 40 ng gDNA, and 8 ng circDNA were used to detect SINV derived vDNA by end point PCR. Mitochondrial DNA was detected in all sample, with circDNA only showing mild enrichment of it, likely due to the limited efficiency of the plasmid prep kit (used for circDNA isolation) in extracting genetic material from mitochondria. **B.** Distribution of transposon-mapping reads in SINV-transposon (TE) hybrid paired reads. **C.** Hybrid paired-end reads between CFAV and Ty1/copia element 56. Viral-mapping (blue dashes, i) and transposon-mapping (red dashes, ii) paired read sequences are shown on the same line (across both virus and transposon). Reads are ordered based on the alignment to the CFAV viral genome starting with the most 5’-end sequence. “Junction” reads refer to any paired-end reads where at least one read contained both virus and transposon sequence. **D.** Coverage of non-hybrid paired-end reads corresponding to (i) Ty3/gypsy or (ii) DENV (ordered vertically by most 5’-end sequence). The respective genome lengths of Ty3/gypsy Ele73 and DENV are 5,131 and 10,700 nt. **E.** Absolute quantification of SINV RNA levels and of two selected host mRNAs found (AAEL011197) or not (rpS17) in episomal DNA in SINV infected Aag2 cells.

**Figure 6 - Supplement.**
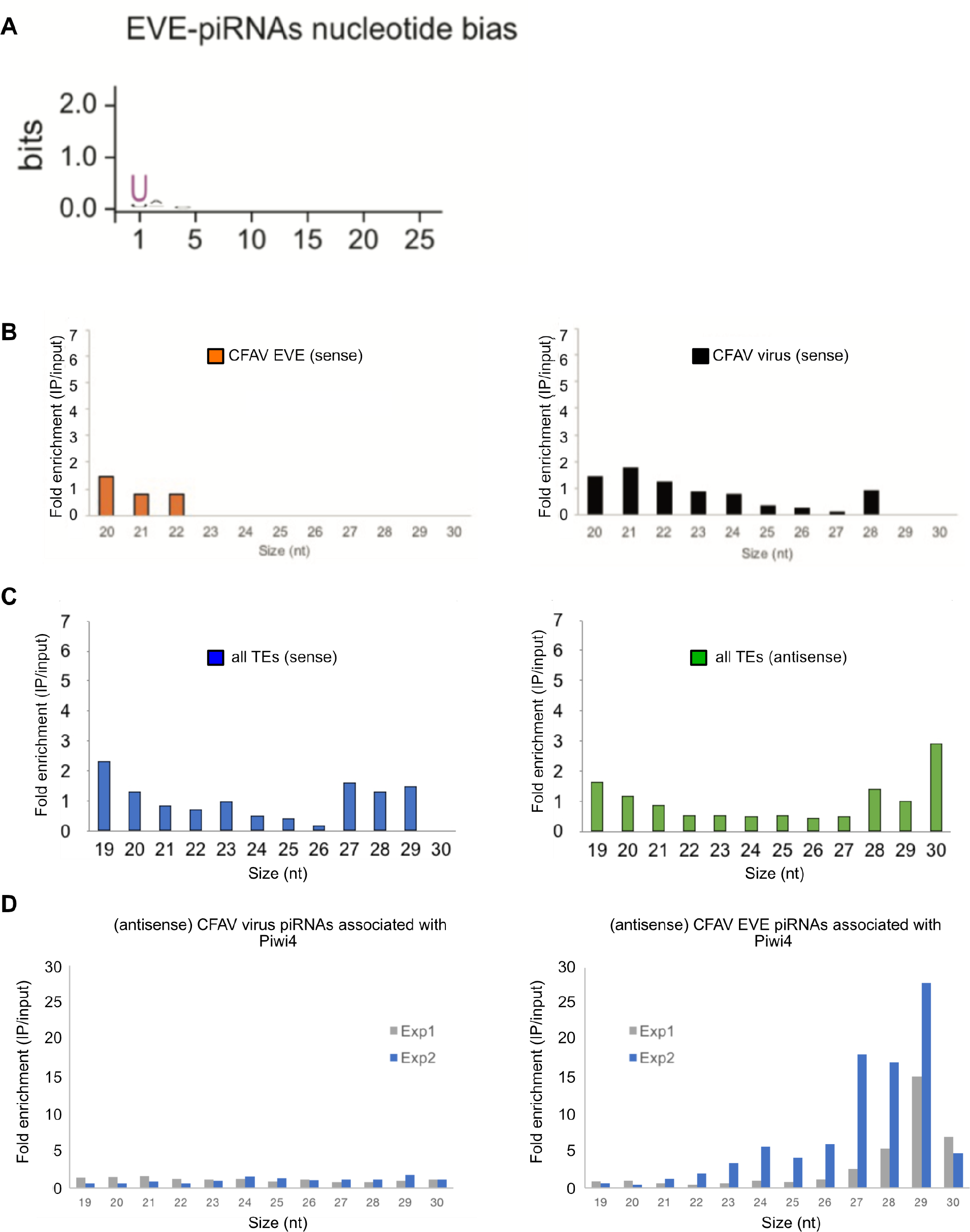
Piwi4 binds to bona-fide antisense v-piRNAs produced by genomic CFAV-EVEs. **A.** The base bias for each position of EVE-derived piRNAs co-immunoprecipitated with Piwi4-FLAG shows a Uracil bias at position 1, characteristic of antisense Piwi-associated piRNAs (shown in bits). **B.** Lack of enrichment of sense CFAV EVE (left panel) or virus (right panel)-derived small RNA in Piwi4-IP fraction compared to input sample. **C.** Lack of enrichment of sense (left panel) or antisense (right panel) TE-targeting piRNAs in Piwi4-IP fraction compared to input sample. **D.** Confirmation of specific enrichment for CFAV EVE-derived antisense piRNAs Piwi4-IP fraction compared to input sample in two additional independent experiments (Exp1 and Exp2).

**Figure 7 - Supplement.**
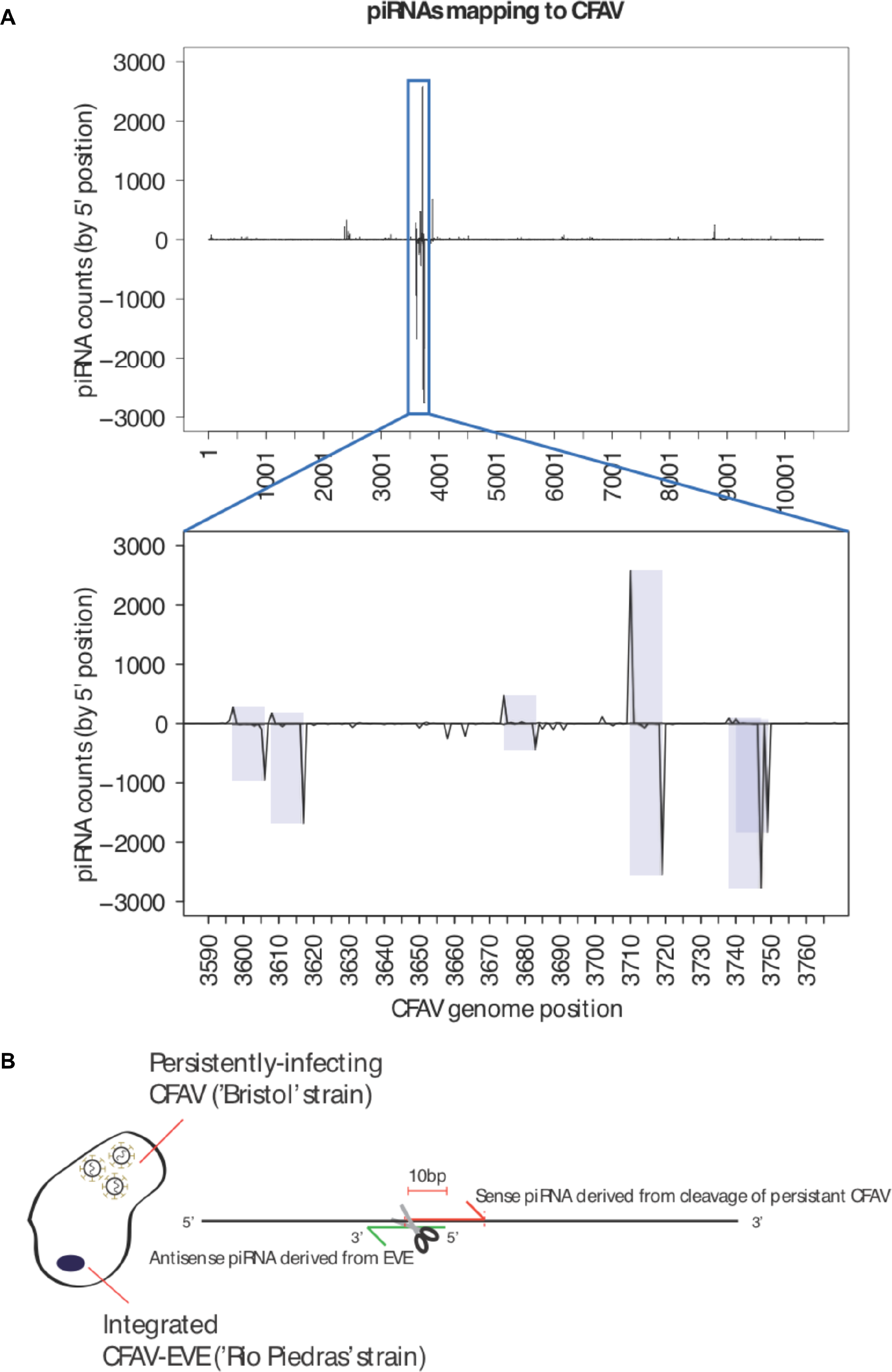
piRNA mapping to CFAV viral genome and EVEs reveals multiple signatures of CFAV virus targeting by EVE-derived piRNAs. **A.** Identification of multiple sense/antisense CFAV piRNAs pairs showing typical ping-pong, 10 base offset signatures. **B.** Schematic of the relationship between an antisense CFAV EVE derived piRNA and its sense CFAV virus piRNA counterpart.

## References

1. Blair CD, Olson KE. The Role of RNA Interference (RNAi) in Arbovirus-Vector Interactions. Viruses. 2015;7:820–843. doi:10.3390/v7020820

2. van Rij RP, Saleh M-C, Berry B, Foo C, Houk A, Antoniewski C, et al. The RNA silencing endonuclease Argonaute 2 mediates specific antiviral immunity in Drosophila melanogaster. Genes Dev. 2006;20:2985–2995. doi:10.1101/gad.1482006

3. Scott JC, Brackney DE, Campbell CL, Bondu-Hawkins V, Hjelle B, Ebel GD, et al. Comparison of Dengue Virus Type 2-Specific Small RNAs from RNA Interference-Competent and -Incompetent Mosquito Cells. PLoS Negl Trop Dis. 2010;4. doi:10.1371/journal.pntd.0000848

4. Hess AM, Prasad AN, Ptitsyn A, Ebel GD, Olson KE, Barbacioru C, et al. Small RNA profiling of Dengue virus-mosquito interactions implicates the PIWI RNA pathway in anti-viral defense. BMC Microbiol. 2011;11:45. doi:10.1186/1471-2180-11-45

5. Morazzani EM, Wiley MR, Murreddu MG, Adelman ZN, Myles KM. Production of Virus-Derived Ping-Pong-Dependent piRNA-like Small RNAs in the Mosquito Soma. PLoS Pathog. 2012;8. doi:10.1371/journal.ppat.1002470

6. Vodovar N, Bronkhorst AW, van Cleef KWR, Miesen P, Blanc H, van Rij RP, et al. Arbovirus-Derived piRNAs Exhibit a Ping-Pong Signature in Mosquito Cells. PLoS One. 2012;7. doi:10.1371/journal.pone.0030861

7. Goic B, Stapleford KA, Frangeul L, Doucet AJ, Gausson V, Blanc H, et al. Virus-derived DNA drives mosquito vector tolerance to arboviral infection. Nat Commun. 2016;7. doi:10.1038/ncomms12410

8. Miesen P, Girardi E, van Rij RP. Distinct sets of PIWI proteins produce arbovirus and transposon-derived piRNAs in Aedes aegypti mosquito cells. Nucleic Acids Res. 2015;43:6545–6556. doi:10.1093/nar/gkv590

9. Varjak M, Maringer K, Watson M, Sreenu VB, Fredericks AC, Pondeville E, et al. Aedes aegypti Piwi4 Is a Noncanonical PIWI Protein Involved in Antiviral Responses. mSphere. 2017;2. doi:10.1128/mSphere.00144-17

10. Brennecke J, Aravin AA, Stark A, Dus M, Kellis M, Sachidanandam R, et al. Discrete small RNA-generating loci as master regulators of transposon activity in Drosophila. Cell. 2007;128:1089–1103. doi:10.1016/j.cell.2007.01.043

11. Petit M, Mongelli V, Frangeul L, Blanc H, Jiggins F, Saleh M-C. piRNA pathway is not required for antiviral defense in Drosophila melanogaster. PNAS. 2016; 201607952. doi:10.1073/pnas.1607952113

12. Arensburger P, Hice RH, Wright JA, Craig NL, Atkinson PW. The mosquito Aedes aegypti has a large genome size and high transposable element load but contains a low proportion of transposon-specific piRNAs. BMC Genomics. 2011;12:606. doi:10.1186/1471-2164-12-606

13. Lewis SH, Salmela H, Obbard DJ. Duplication and Diversification of Dipteran Argonaute Genes, and the Evolutionary Divergence of Piwi and Aubergine. Genome Biol Evol. 2016;8:507–518. doi:10.1093/gbe/evw018

14. Varjak M, Dietrich I, Sreenu VB, Till BE, Merits A, Kohl A, et al. Spindle-E Acts Antivirally Against Alphaviruses in Mosquito Cells. Viruses. 2018;10. doi:10.3390/v10020088

15. Tassetto M, Kunitomi M, Andino R. Circulating Immune Cells Mediate a Systemic RNAi-Based Adaptive Antiviral Response in Drosophila. Cell. 2017;169:314–325.e13. doi:10.1016/j.cell.2017.03.033

16. Holmes EC. The Evolution of Endogenous Viral Elements. Cell Host & Microbe. 2011;10:368–377. doi:10.1016/j.chom.2011.09.002

17. Palatini U, Miesen P, Carballar-Lejarazu R, Ometto L, Rizzo E, Tu Z, et al. Comparative genomics shows that viral integrations are abundant and express piRNAs in the arboviral vectors Aedes aegypti and Aedes albopictus. BMC Genomics. 2017;18:512. doi:10.1186/s12864-017-3903-3

18. Whitfield ZJ, Dolan PT, Kunitomi M, Tassetto M, Seetin MG, Oh S, et al. The Diversity, Structure, and Function of Heritable Adaptive Immunity Sequences in the Aedes aegypti Genome. Curr Biol. 2017;27:3511–3519.e7. doi:10.1016/j.cub.2017.09.067

19. Schnettler E, Donald CL, Human S, Watson M, Siu RWC, McFarlane M, et al. Knockdown of piRNA pathway proteins results in enhanced Semliki Forest virus production in mosquito cells. J Gen Virol. 2013;94:1680–1689. doi:10.1099/vir.0.053850-0

20. Dietrich I, Jansen S, Fall G, Lorenzen S, Rudolf M, Huber K, et al. RNA Interference Restricts Rift Valley Fever Virus in Multiple Insect Systems. mSphere. 2017;2. doi:10.1128/mSphere.00090-17

21. Varjak M, Donald CL, Mottram TJ, Sreenu VB, Merits A, Maringer K, et al. Characterization of the Zika virus induced small RNA response in Aedes aegypti cells. PLoS Negl Trop Dis. 2017;11:e0006010. doi:10.1371/journal.pntd.0006010

22. Horwich MD, Li C, Matranga C, Vagin V, Farley G, Wang P, et al. The Drosophila RNA methyltransferase, DmHen1, modifies germline piRNAs and single-stranded siRNAs in RISC. Curr Biol. 2007;17:1265–1272. doi:10.1016/j.cub.2007.06.030

23. Goic B, Vodovar N, Mondotte JA, Monot C, Frangeul L, Blanc H, et al. RNA-mediated interference and reverse transcription control the persistence of RNA viruses in the insect model Drosophila. Nat Immunol. 2013;14:396–403. doi:10.1038/ni.2542

24. Nag DK, Brecher M, Kramer LD. DNA forms of arboviral RNA genomes are generated following infection in mosquito cell cultures. Virology. 2016;498:164–171. doi:10.1016/j.virol.2016.08.022

25. Hotta Y, Bassel A. MOLECULAR SIZE AND CIRCULARITY OF DNA IN CELLS OF MAMMALS AND HIGHER PLANTS. Proc Natl Acad Sci USA. 1965;53:356–362.

26. Jones RS, Potter SS. L1 sequences in HeLa extrachromosomal circular DNA: evidence for circularization by homologous recombination. Proc Natl Acad Sci USA. 1985;82:1989–1993.

27. Flavell AJ, Ish-Horowicz D. Extrachromosomal circular copies of the eukaryotic transposable element copia in cultured Drosophila cells. Nature. 1981;292:591–595.

28. Møller HD, Bojsen RK, Tachibana C, Parsons L, Botstein D, Regenberg B. Genome-wide Purification of Extrachromosomal Circular DNA from Eukaryotic Cells. J Vis Exp. 2016; e54239 |. doi:10.3791/54239

29. Stollar V, Thomas VL. An agent in the Aedes aegypti cell line (Peleg) which causes fusion of Aedes albopictus cells. Virology. 1975;64:367–377.

30. Lai MM. Genetic recombination in RNA viruses. Curr Top Microbiol Immunol. 1992;176:21–32.

31. Skala AM. Retroviral DNA Transposition: Themes and Variations. Microbiol Spectr. 2014;2. doi:10.1128/microbiolspec.MDNA3-0005-2014

32. Zhang J, Tang L-Y, Li T, Ma Y, Sapp CM. Most Retroviral Recombinations Occur during Minus-Strand DNA Synthesis. J Virol. 2000;74:2313–2322.

33. Horie M, Honda T, Suzuki Y, Kobayashi Y, Daito T, Oshida T, et al. Endogenous non-retroviral RNA virus elements in mammalian genomes. Nature. 2010;463:84–87. doi:10.1038/nature08695

34. Fort P, Albertini A, Van-Hua A, Berthomieu A, Roche S, Delsuc F, et al. Fossil rhabdoviral sequences integrated into arthropod genomes: ontogeny, evolution, and potential functionality. Mol Biol Evol. 2012;29:381–390. doi:10.1093/molbev/msr226

35. Taylor DJ, Leach RW, Bruenn J. Filoviruses are ancient and integrated into mammalian genomes. BMC Evol Biol. 2010;10:193. doi:10.1186/1471-2148-10-193

36. Crochu S, Cook S, Attoui H, Charrel RN, De Chesse R, Belhouchet M, et al. Sequences of flavivirus-related RNA viruses persist in DNA form integrated in the genome of Aedes spp. mosquitoes. J Gen Virol. 2004;85:1971–1980. doi:10.1099/vir.0.79850-0

37. Katzourakis A, Gifford RJ. Endogenous viral elements in animal genomes. PLoS Genet. 2010;6:e1001191. doi:10.1371/journal.pgen.1001191

38. Bushman FD. Targeting survival: integration site selection by retroviruses and LTR-retrotransposons. Cell. 2003;115:135–138.

39. Peleg J. Growth of arboviruses in primary tissue culture of Aedes aegypti embryos. Am J Trop Med Hyg. 1968;17:219–223.

40. Olson KE, Bonizzoni M. Nonretroviral integrated RNA viruses in arthropod vectors: an occasional event or something more? Curr Opin Insect Sci. 2017;22:45–53. doi:10.1016/j.cois.2017.05.010

41. Marklewitz M, Zirkel F, Kurth A, Drosten C, Junglen S. Evolutionary and phenotypic analysis of live virus isolates suggests arthropod origin of a pathogenic RNA virus family. Proc Natl Acad Sci U S A. 2015;112:7536–7541. doi:10.1073/pnas.1502036112

42. Hall RA, Bielefeldt-Ohmann H, McLean BJ, O’Brien CA, Colmant AMG, Piyasena TBH, et al. Commensal Viruses of Mosquitoes: Host Restriction, Transmission, and Interaction with Arboviral Pathogens. Evol Bioinform Online. 2017;12:35–44. doi:10.4137/EBO.S40740

43. de Vanssay A, Bougé A-L, Boivin A, Hermant C, Teysset L, Delmarre V, et al. Paramutation in Drosophila linked to emergence of a piRNA-producing locus. Nature. 2012;490:112–115. doi:10.1038/nature11416

44. Sladowski D, Steer SJ, Clothier RH, Balls M. An improved MTT assay. J Immunol Methods. 1993;157:203–207.

45. Takeuchi H, Baba M, Shigeta S. An application of tetrazolium (MTT) colorimetric assay for the screening of anti-herpes simplex virus compounds. J Virol Methods. 1991;33:61–71.

46. Anderson MAE, Gross TL, Myles KM, Adelman ZN. Validation of novel promoter sequences derived from two endogenous ubiquitin genes in transgenic Aedes aegypti. Insect Mol Biol. 2010;19:441–449. doi:10.1111/j.1365-2583.2010.01005.x

47. Tong H, Schliekelman P, Mrázek J. Unsupervised statistical discovery of spaced motifs in prokaryotic genomes. BMC Genomics. 2017;18. doi:10.1186/s12864-016-3400-0

48. Chen C, Khaleel SS, Huang H, Wu CH. Software for pre-processing Illumina next-generation sequencing short read sequences. Source Code Biol Med. 2014;9:8. doi:10.1186/1751-0473-9-8

49. Langmead B, Trapnell C, Pop M, Salzberg SL. Ultrafast and memory-efficient alignment of short DNA sequences to the human genome. Genome Biol. 2009;10:R25. doi:10.1186/gb-2009-10-3-r25

50. Watson M, Schnettler E, Kohl A. viRome: an R package for the visualization and analysis of viral small RNA sequence datasets. Bioinformatics. 2013;29:1902–1903. doi:10.1093/bioinformatics/btt297

51. Crooks GE, Hon G, Chandonia J-M, Brenner SE. WebLogo: A Sequence Logo Generator. Genome Res. 2004;14:1188–1190. doi:10.1101/gr.849004

52. Patro R, Duggal G, Love MI, Irizarry RA, Kingsford C. Salmon provides fast and bias-aware quantification of transcript expression. Nat Methods. 2017;14:417–419. doi:10.1038/nmeth.4197

